# Neural knowledge assembly in humans and deep networks

**DOI:** 10.1101/2021.10.21.465374

**Authors:** Stephanie Nelli, Lukas Braun, Tsvetomira Dumbalska, Andrew Saxe, Christopher Summerfield

## Abstract

Human understanding of the world can change rapidly when new information comes to light, such as when a plot twist occurs in a work of fiction. This flexible “knowledge assembly” requires few-shot reorganisation of neural codes for relations among objects and events. However, existing computational theories are largely silent about how this could occur. Here, participants learned a transitive ordering among novel objects within two distinct contexts, before exposure to new knowledge that revealed how they were linked. BOLD signals in dorsal frontoparietal cortical areas revealed that objects were rapidly and dramatically rearranged on the neural manifold after minimal exposure to linking information. We then adapt stochastic online gradient descent to permit similar rapid knowledge assembly in a neural network model.

## Introduction

To make sense of the world, we need to know how objects, people and places relate to one another. Understanding how relational knowledge is acquired, organised and used for inference has become a frontier topic in both neuroscience and machine learning research [1-7]. Since Tolman, neuroscientists have proposed that when ensembles of states are repeatedly co-experienced, they are mentally organised into cognitive maps whose geometry mirrors the external environment [8-12]. On a neural level, the associative distance between objects or locations (i.e. how related they are in space or time), has been found to covary with similarity (or dissimilarity) among neural coding patterns. Some neural signals, especially in medial temporal lobe structures, may even explicitly encode relational information about how space is structured or how knowledge hierarchies are organised [13-15].

A striking aspect of cognition is that these knowledge structures can be rapidly reconfigured when new information becomes available. For example, a plot twist in a film might require the viewer to rapidly and dramatically reconsider a protagonist’s motives, or an etymylogical insight might allow a reader to suddenly understand the connection between two words. Here, we dub this process “knowledge assembly” because it requires existing knowledge to be rapidly (re-)assembled on the basis of minimal new information. How do brains update knowledge structures, selectively updating certain relations while keeping others in tact? In machine learning research [16,17], solutions to the general problem of building rich conceptual knowledge structures include graph-based architectures [18], modular networks [19], probabilistic programs [20], and deep generative models [21]. However, whilst these artificial tools can allow for expressive mental representation or powerful inference, they tend to learn slowly and require dense supervision, making them implausibile models of knowledge assembly and limiting their scope as theories of biological learning.

How, then, does knowledge assembly occur in humans? We designed a task in which human participants learned the rank of adjacent items within two ordered sets of novel objects occurring in distinct temporal contexts. Participants acquired and generalised the transitive relations both within and between contexts, and did so in a fashion qualitatively identical to a feedforward neural network. The geometry of multivoxel BOLD signals recorded from dorsal stream structures suggested that humans solved the task by representing objects on two parallel mental lines, one for each context, building on previous findings [22-25]. This coding strategy mirrored that observed in the hidden layer of the neural network. We then provided a very small number of ‘list linking’ training examples meant to imply that the two ordered sets in fact lay on a single continuum. Our participants rapidly infered the full set of resulting transitive relations given this minimal (and potentially ambiguous) information, as found previously in humans [26] and macaques [27]. As we recorded human BOLD signals both before and after this brief training period, we observed remarkable few-shot adjustments in neural geometries consequent from the new information. We then describe a theory of how knowledge can be rapidly assembled using a version of the artificial neural network model, providing a computational account of the behavioural and neural results observed in humans.

## Results

Human participants (n = 34) performed a computerised task that involved making decisions about novel visual objects. Each object *i* was randomly assigned a ground truth rank (*i*_1_ − *i*_12_) on the nonsense dimension of “brispiness” (Fig. 1A; see Methods; where *i*_1_ is the most brispy and *i*_12_ is the least). During initial training (train_short), the 12 objects were split into two distinct sets (items *i*_1_ − *i*_6_ and *i*_7_ − *i*_12_) and presented in alternating blocks (contexts; see Methods). Within each context, participants were asked to indicate with a button press which of two objects with adjacent rank (e.g. *i*_3_ and *i*_4_) as more (or less) “brispy”, receiving fully informative feedback (Fig. 1B, upper panel). Note that this training regime allowed participants to infer ranks within a set (i.e. within *i*_1_ − *i*_6_ and *i*_7_ − *i*_12_) but betrayed no information about the ground truth relation between the two sets (e.g. *i*_2_ < *i*_9_). Participants were trained on adjacent relations to a predetermined criterion, with final training accuracy reaching 95.6 ± 2.9% (mean ± SD; Fig. 1C; see Methods). The use of novel objects [28] and a nonword label was designed to minimise participants’ tendency to use prior information when solving the task.

**Figure 1.**
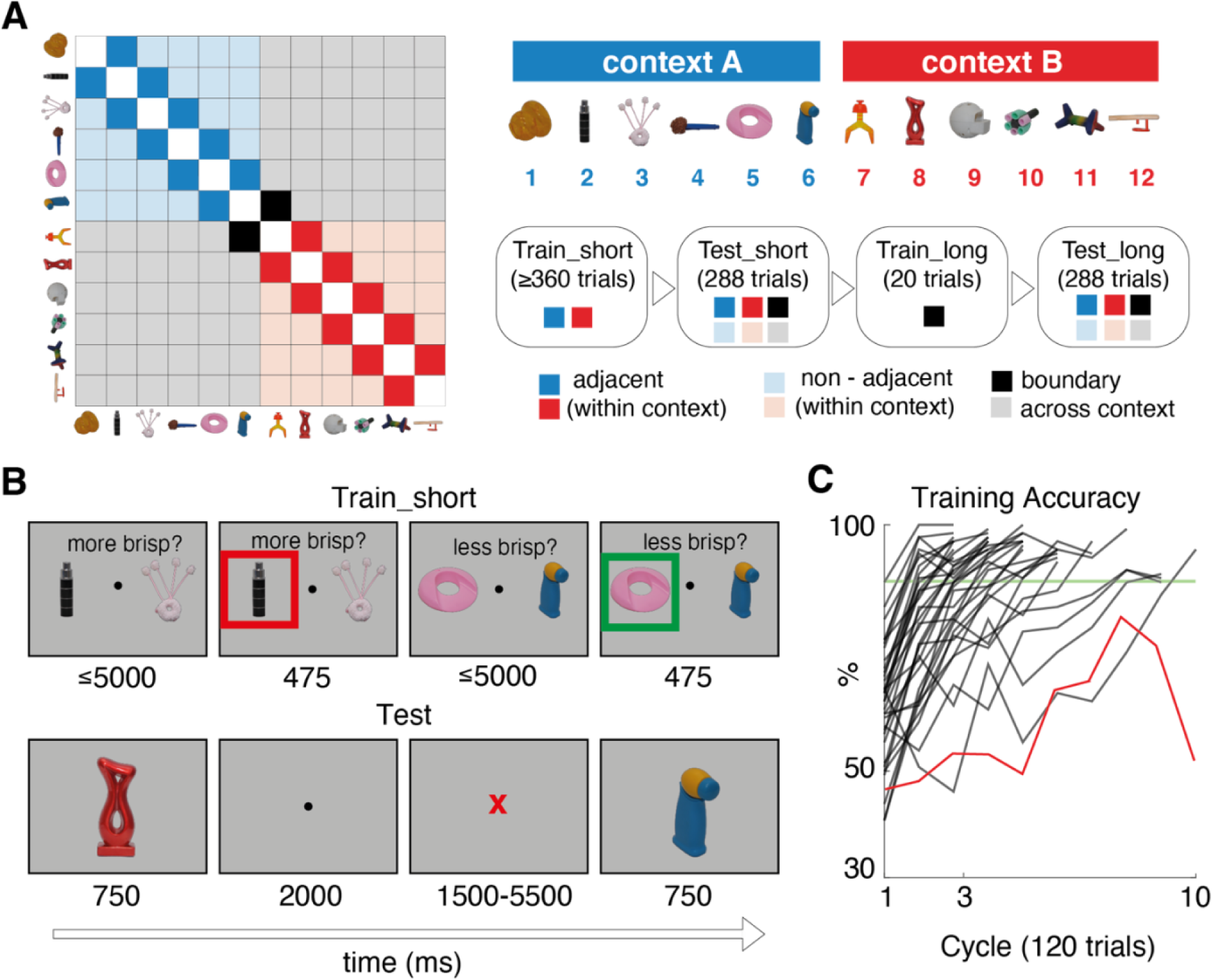
Task and design. A. Left: matrix illustrating training and testing conditions for an example set of objects ordered by rank (on x and y axes). Each entry indicates a pair of stimuli defined by their row and column. Colours signal when the pair was trained or tested. For example, dark blue and red squares are within-context pairs, shown during train_short. In addition to these, lighter blue and red squares (trained non-adjacent) and grey (untrained) are pairs tested during test_short. The black squares are the pairs shown during boundary training (train_long). Right: schematic of experimental sequence and legend. Although we use the same set of objects for display purposes, note that each participant viewed a randomly sampled set of novel objects B. Example trial sequence during training (upper) and test (lower). Numbers below each example screen show the frame duration in ms. Timings for test were chosen to assist with BOLD modelling. C. Percentage accuracy over blocks during training for each individual. Stopping criterion is shown as a green line. The excluded participant is shown as a red trace. A training “cycle” consists of two blocks (one for each set of items). More details are provided in the Online Methods.

After training, participants entered the scanner and performed a first test phase [test_short] in which they viewed objects one by one that were sampled randomly from across the full range (*i*_1_ − *i*_12_). The task required them to report the “brispiness” of each object relative to its predecessor (i.e. a 1-back task) with a button press (Fig. 1B, lower panel). Therefore, the test phase involved comparisons of trained (adjacent) pairs within context (e.g. *i*_3_ and *i*_4_), untrained (non-adjacent) pairs within context (e.g. *i*_3_ and *i*_6_) and untrained pairs across contexts (e.g. *i*_3_ and *i*_10_). Importantly, participants did not receive trialwise feedback on their choices during the test phase.

Our first question was whether humans generalised knowledge about object *brispiness* both within and between contexts during the test_short phase. We collapse across the two contexts as there was no difference in either reaction times (RT) or accuracy between them (both p-values > 0.3). Participants performed above chance both on adjacent pairs on which they had been trained (e.g. *i*_3_ and *i*_4_ or *i*_9_ and *i*_10_) [mean accuracy = 86.0 ± 10.4, t-test against 50%, t_33_ = 20.5, p < 0.001] but also on untrained, nonadjacent pairs for which transitive inference was required (e.g. *i*_3_ and *i*_6_ or *i*_7_ and *i*_10_) (Fig. 2A) [mean accuracy 96.7 ± 19.2, t_33_ = 83.7, p < 0.001]. In fact, they were faster and more accurate for comparisons between non-adjacent than adjacent items (Fig. 2B) [accuracy: t_33_ = 7.8, p < 0.001; RT: t_33_ = 11.7, p < 0.001]. This was driven by an increase in accuracy (and decrease in RT) with growing distance between comparanda (Fig. 2A-B, right panels) [accuracy: *β* = 3.4% per rank; t_33_ = 7.7, p < 0.001. RT: = 72 ms faster per rank; t_33_ = -7.8, p < 0.001; *βs* obtained with a regression model], known as the “symbolic distance” effect [26,29]. We note that this result is not readily explained by models of transitive inference based on spreading pairwise associations through recurrent dynamics [30].

**Fig 2.**
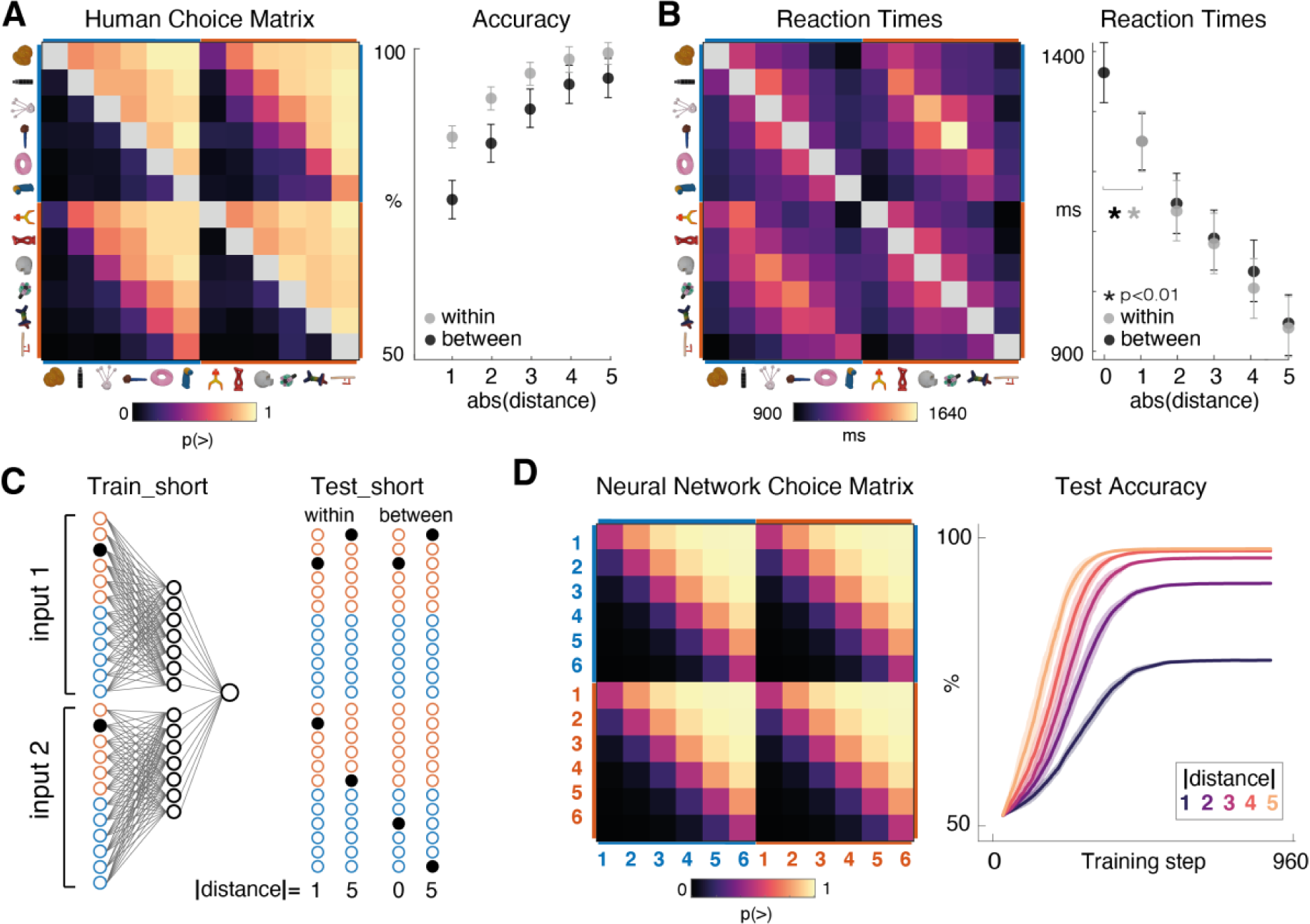
Behaviour in humans and neural networks. A. Left panel: human choice matrix. The colour of each entry indicates the probability of responding “greater than” during test_short for the pair of items defined by the row and column. Colour scale is shown below the plot. Object identities are shown for illustration only (and were in fact resampled for each participant). Right panel: accuracy as a function of symbolic distance shown separately for within-context (e.g. *i*_3_ and *i*_5_; grey dots) and between context (e.g. *i*_3_ and *i*_9_; black dots) judgements. For between, accuracy data are with respect to a ground truth in which ranks are perfectly generalised across contexts (e.g. they infer that g. *i*_2_ > *i*_9_;). Errors bars are S.E.M. B. Equivalent data for reaction times. Note that a symbolic distance of zero was possible across contexts (e.g. *i*_2_ vs. *i*_8_) for which there was no correct answer but an RT was measurable. C. Left panel: neural network architecture and training scheme. Input nodes are colored red and blue to denote the relevant context. Black dots illustrate an example training train in which objects *i*_2_ and *i*_3_ are shown. Right panel: example test trials both within and across context, with the symbolic distance signalled below. D. Left Panel: Choice matrix for the neural network, in the same format as A. Right Panel: Learning curves (showing accuracy over training epochs) for the neural network, shown separately for trials with different levels of symbolic distance. Shading is 1 S.E.M over network replicants. Note that like humans, despite being trained exclusively on adjacent items, neural networks learned faster and performed better on non-adjacent items.

Moreover, behaviour also indicated *how* participants compared ranks between contexts before ground truth was revealed. For example, they tended to infer *i*_7_ > *i*_2_ and *i*_4_ > *i*_11_ (Fig 2A). This implies a natural tendency to match rank orderings between contexts (e.g. that the 3^rd^ item in one set was ranked higher than the 4^th^ in the other) in the absence of information about how objects were related across contexts. In line with this, we quantified between-context accuracy relative to an agent that generalises perfectly between contexts and found that between-context accuracy was well above chance for adjacent [75.9 ± 20.9 mean ± SD; t_33_ = 7.3, p < 0.001] and non-adjacent [91.5 ± 9.3 mean ± SD; t_33_ = 14.2, p < 0.001] trials.

We also observed a *between*-*context* symbolic distance effect in reaction times (Fig 2B) [accuracy: 4.9% per rank, t_33_ = 7.5, p < 0.001. RT: beta = 75 ms faster per rank; t_33_ = -7.8, p < 0.001]. Participants were slowest when comparing items with equivalent rank across contexts (e.g. comparing *i*_2_ and *i*_8_, which were both ranked 2^nd^ within their respective contexts), responding more slowly than for adjacent items both within [t_33_ = 3.23, p < 0.004], and between [t_33_ = 3.66, p < 0.001] contexts. Overall, these results are consistent with previous findings in both humans and monkeys, and have been taken to imply that participants automatically infer and represent the ordinal position of each item in the set [31].

Next, to understand the computational underpinnings of this behaviour and neural coding, we trained a neural network to solve an equivalent transitive inference problem. The network had a two-layer feedforward architecture with symmetric input weights, and was trained in a supervised fashion using stochastic online gradient descent [SGD] (Fig. 2C). For this modelling exercise, we replaced the unrelated object images seen by participants for orthogonal (one-hot) vector inputs. On each trial the network received two inputs, denoting the images shown on the right and left of the screen, and (just like participants) was required to output whether one was “more” or “less” than the other (see Methods). At the point at which we terminated training (960 total trials, or 8 blocks of 60 trials each per context; see Methods), the network reached an average test accuracy of 97.74% on unseen comparisons (between nonadjacent pairs) and 79.04% for adjacent (trained) pairs (Fig. 2D, right panel). It showed a qualitatively identical pattern of generalisation within and between contexts, such that accuracy grew with rank distance (Fig. 2D, left panel). Choice matrices for the humans and neural networks were highly correlated [*r* = 0.98, p < 0.001 for averaged choice matrices; single participants *r* = 0.82 ± 0.13 mean ± SD, all p-values < 0.001].

After training, we examined neural geometry in the neural network by probing it with each (single) item *i*_1_ − *i*_12_ in turn and calculating a representational dissimilarity matrix (RDM) from resultant hidden layer activations (Fig. 3A top row). We then used multidimensional scaling (MDS) to visualise the similarity structure in just two dimensions (Fig. 3A bottom row). As training progressed, the network learned to represent the items in order of brispiness along two adjacent parallel neural lines. We know from recent work that a low dimensional solution is only guaranteed when the hidden layer weights are initialised from very small values, sometimes known as the “rich” training regime [32]. After training in this regime, the Pearson’s correlation between the data RDM from the hidden layer of the neural network (RDM_NN_) and an idealised distance matrix for parallel lines (RDM_mag_; Fig. 3B top panel) was > 0.99 for all networks trained (p < 0.001 for each of 20 networks). However, we observed no correlation between RDM_NN_ and an RDM coding for distance between contexts (RDM_ctx_; Fig. 3B bottom panel) [Pearson *r* ≤ 0.1, p > 0.4 for all cases], consistent with the observation that the magnitude lines were not just adjacent but fully overlapping by the end of training (Fig. 3A).

**Fig 3.**
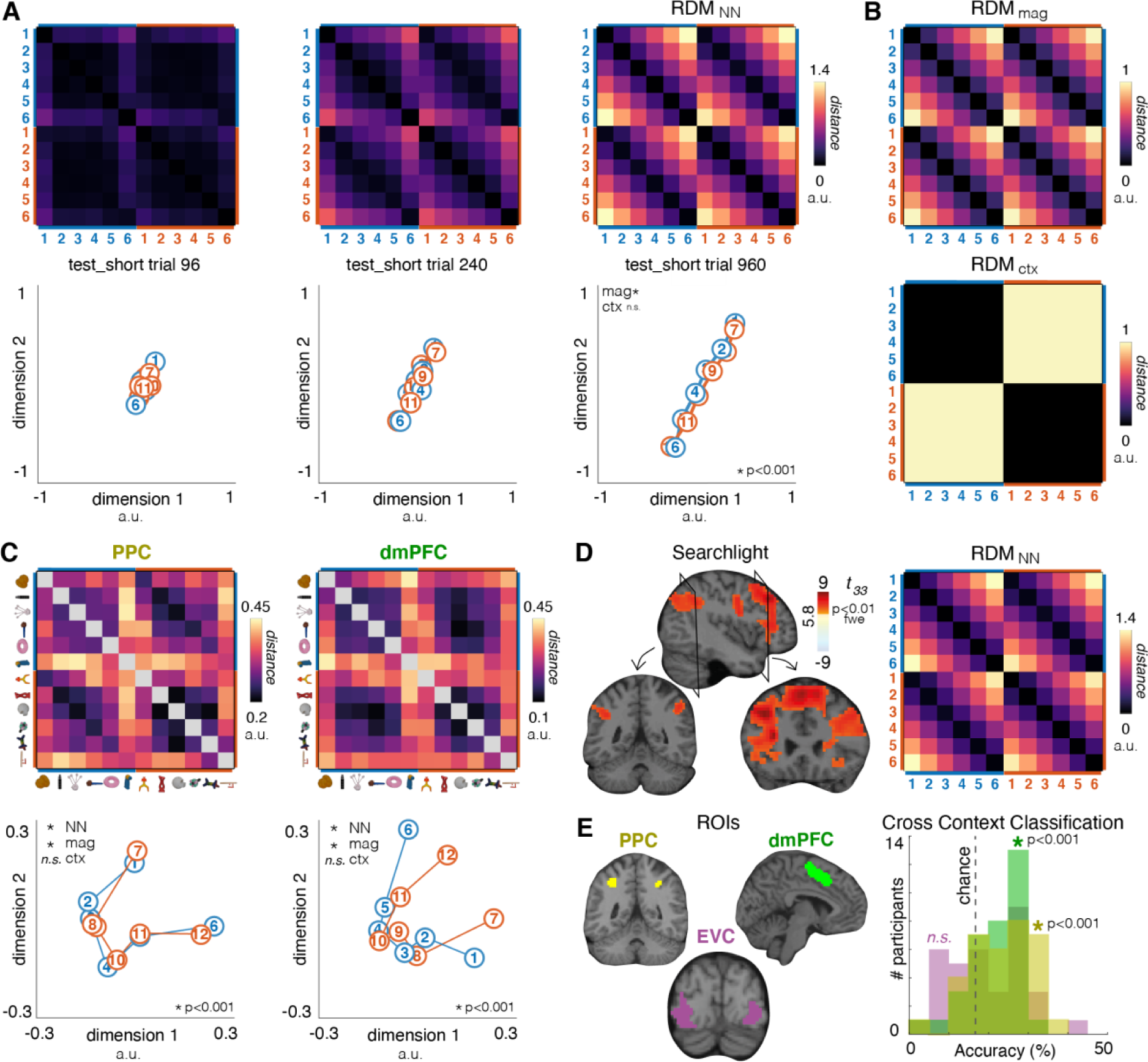
Data from artificial networks and human BOLD signals. A. Upper panels: RDM for the neural network. Each entry shows the distance between hidden unit activations evoked by a pair of stimuli, for three example timepoints during training. Lower panels: MDS plot in 2D for the RDM above. Each circle is a stimulus, coloured by its context. Distances between circles conserve similarities in the RDM. Note the emergence of two parallel lines. B. Model RDMs for magnitude (assumes linear spacing between ranks) and context (assumes a fixed distance between contexts). C. Upper panels: neural data RDMs from patterns of BOLD in the PPC (left) and dmPFC (right) regions of interest (ROIs). Lower panels: 2D MDS on BOLD data. Red and blue lines denote the two contexts; numbers circles denote items, with their rank signalled by the inset number. D. Voxels correlating reliably with the terminal RDM from the neural network (RDM_NN_, see right panel) rendered onto saggital (upper) and coronal (lower) slices of a standardised brain, at a threshold of FWE p < 0.01. E. Left panel: Regions of interest (ROIs) in posterior parietal cortex (PPC, yellow), dorsomedial prefrontal cortex (PFC, green) and a control region in Visual Cortex (VC, purple). Right panel: Frequencies of accuracy bins over participants for support vector machines (SVMs) trained to distinguish item ranks in one context after training on the other. Three histograms are overlaid, one for each ROI; colours correspond to those for panel D. Dashed line shows chance (16.6%).

Next, we compared the representational geometry observed in the neural network to that recorded in BOLD signals whilst human participants judged the “brispiness” of successive items in the test_short phase. We initially focus on regions of interest (ROIs) derived from an independent task in which participants judged the magnitude of Arabic digits, localised to the posterior parietal cortex (PPC) and dorsomedial prefrontal cortex (dmPFC; see Fig. S1), and later show the involvement of a larger fronto-parietal network using a whole-brain searchlight approach. In both ROIs, we saw strong correlation between the neural data RDM and RDM_mag_ (Fig. 3C) [PPC: t_33_ = 4.2, p < 0.001; dmPFC: t_33_ = 5.7, p < 0.001] but no effect of RDM_ctx_ [t_33_ < 1, p > 0.65 for both regions]. This echoes the data from the hidden layer of the neural network (see Fig. 3A), and accordingly we observed significant correlation with RDM_NN_ in both regions [PPC: t_33_ = 5.3; dmPFC: t_33_ = 7.2, both p < 0.001]. These effects all still held when we defined similarity across (rather than within) scanner runs using a crossvalidated RSA approach [RDM_mag_: PPC: t_33_ = 4.8, p < 0.001; dmPFC: t_33_ = 4.7, p < 0.001; RDM_ctx_: PPC: t_33_ = 0.3, p = 0.78; dmPFC: t_33_ = 0.4, p = 0.69; RDM_NN_: PPC: t_33_ = 5.4, dmPFC: t_33_ = 6.0, p < 0.001; see Fig. S2].

We visualised the neural geometry of the BOLD signals in both regions after reducing to two dimensions with MDS. In both ROIs, this yielded overlapping neural lines that reflected the rank-order of the novel objects (Fig. 3C, bottom row; see Fig. S3). Restricting our analysis to distances between consecutive objects, we found neural distances involving the end anchors (e.g. *i*_1_ and *i*_6_) tended to be larger than those involving intermediate ranks [*t_33_* = 4.8, p < 0.001; *t_33_* = 4.4, p < 0.001 in PPC and dmPFC respectively]. However, unlike in the neural network, manifolds (number lines) obtained from BOLD data were curved. We note that the curvature of these representational manifolds around their midpoint yields approximately orthogonal axes for rank and uncertainty, and that this phenomenon has been previously observed in scalp EEG recordings [25] and in multi-unit activity from area LIP of the macaque [33].

This similarity between representations in the neural network and human BOLD was confirmed by a whole-brain searchlight approach, for which we report only effects that pass a familywise error (FWE) correction level of p < 0.01 with cluster size > 10 voxels (see Methods). This approach revealed a fronto-parietal network in which multivoxel patterns resembled those for the trained neural network (RDM_NN_; Fig. 3D), with peaks in dmPFC [-33 -3 33; t_33_= 9.3, p_uncorr_ < 0.001] and inferior parietal lobe [right: 51 -45 45; t_33_ = 6.94, p_uncorr_ < 0.001; left: -39 -51 36; t_33_ = 6.93, p_uncorr_ < 0.001]. As expected, this was driven by an explicit representation of magnitude distance, as correlations with RDM_mag_ (Fig. 3B, top panel) peaked in the same regions [dmPFC: -33 -3 33; t_33_= 9.4, p<0.001; inferior parietal lobe right: 48 -45 42; t_33_ = 7.01, p_uncorr_ < 0.001; left: -39 -51 36; t_33_ = 7.15, p_uncorr_ < 0.001]. Notably, we did not observe an effect of RDM_ctx_ (Fig. 3B, bottom panel) [no clusters survived FWE correction, *t_33_* < 2.65, uncorrected p > 0.012], indicating that neural representations for similarly-ranked items within each of the two contexts were effectively superimposed, as in the neural network.

This representational format, whereby ranked items are represented on parallel manifolds, lends itself to generalisation across contexts, i.e. between items with distinct identity but equivalent brispiness [23,24]. To test this, we trained a support vector machine on binary classifications among ranks for context A and evaluated it on the (physically dissimilar) objects in context B. We found above-chance classification in PPC and dmPFC (Fig. 3E) [*t_33_* = 4.39, p < 0.001; *t_33_* = 4.01, p < 0.001], but not in an extrastriate visual cortex ROI that also showed significant activation during the independent localiser [*t_33_* = 1.75, p > 0.08]. These analyses not only cross-validated across runs, but also counterbalanced response contingencies, and so are unlikely to be indexing any spurious effect of motor control. Together, these results show that neural patterns indexed a concept of “brispiness” divorced from the physical properties of the objects themselves.

Next, we turned to our central question of how neural representations are reconfigured following a single piece of new information about the overall knowledge structure. After test_short, participants performed a brief “boundary training” session in the scanner in which they learned that object *i*_7_ (the most brispy object in context B) was less brispy then object *i*_6_ (the least brispy object in context A). This information was acquired over just 20 trials in which participants repeatedly judged whether item *i*_6_ or *i*_7_ was “more” or “less” brispy. Following this boundary training, participants performed a new session test_long which was identical in every respect to test_short.

Our main question was whether and how the boundary training reshaped both behaviour and neural coding for the full set of objects. The average choice and RT matrices observed during test_long are shown in Fig. 4A. As can be seen, on aggregate participants used knowledge of relations between items *i*_6_ and *i*_7_ to correctly infer that all objects lay on a single long axis of brispiness (ranked 1-12). We confirmed this in two ways. First, unlike in test_short, items in context A (*i*_1_ − *i*_6_) were mostly ranked as more brispy than items in context B (*i*_7_ − *i*_12_) and the symbolic distance effect now spanned the whole range of items 1-12 (with a “dip” near the boundary between contexts; Fig. 4B, left) [accuracy: 2.1% per rank; t_33_ = 7.8, p < 0.001. RT: - 29 ms per rank; t_33_ = -10.4, p < 0.001]. Next, we directly quantified the full pattern of responses seen in Fig. 4A by constructing idealised ground truth choice and reaction time (RT) matrices (Fig. S4A). These matrices reflected the assumption that the items either lay on two parallel short axes (as most participants inferred in test_short) or a single long axis (as was correct in test_long). Fitting these to human behavioral matrices using competitive regressions, we found that while the long axis matrix fit the test_long behavioral data [Choice: t_33_ = 9.2, p < 0.0001; RTs: t_33_ = 7.9, p < 0.0001] there remained a strong residual fit to the short axis choice and RT patterns [Choice: t_33_ = 5.0, p < 0.0001; RTs: t_33_ = 3.5, p < 0.01].

**Figure 4:**
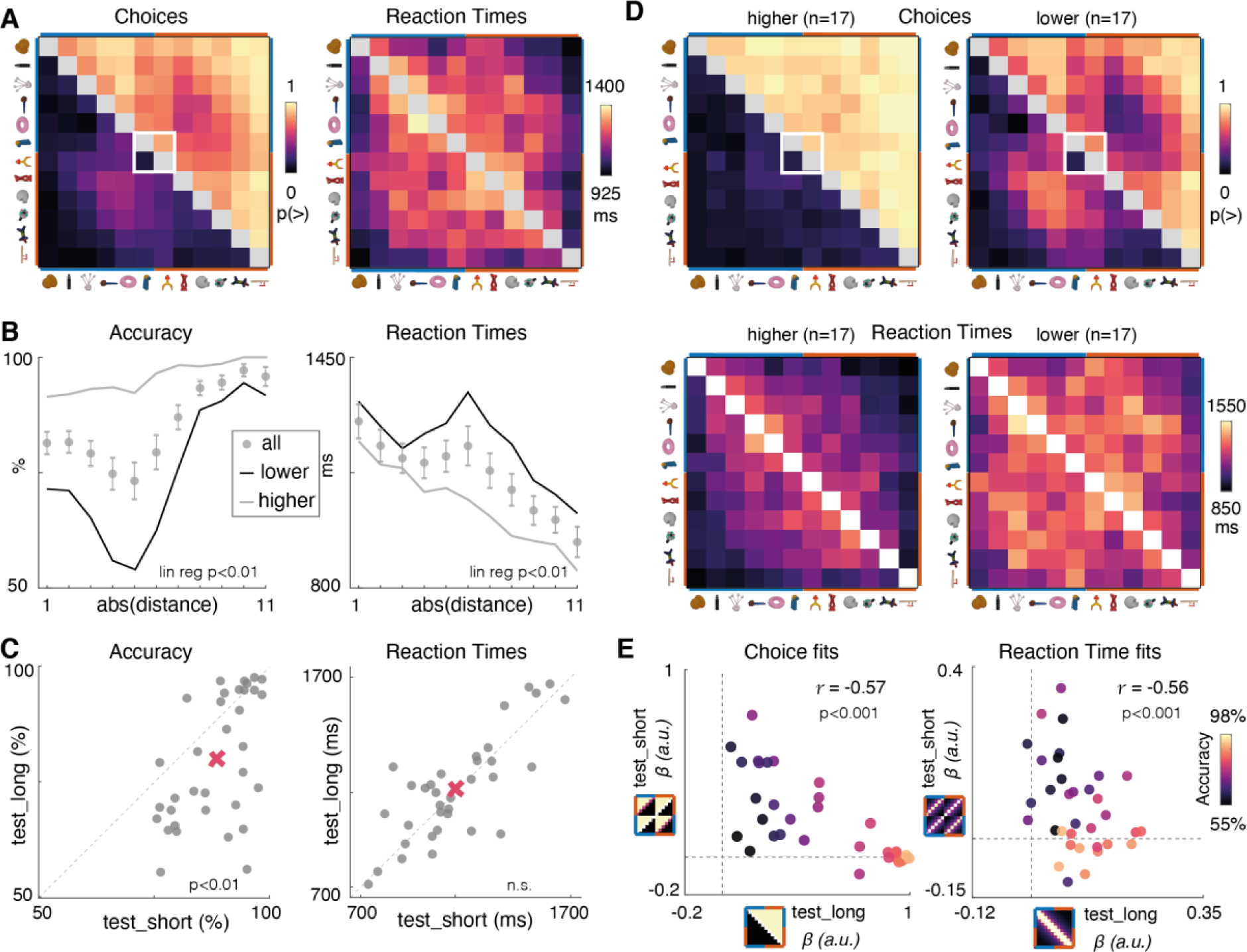
Test_long Behavior. A. Right panel: choice matrices from human participants after boundary training. Format as for Fig. 2A. The white box indicates the two items viewed during boundary training. Note that now, on average choices respect the ground truth rank (*i*_1_ − *i*_12_). Left panel: same for RTs. B. Mean accuracy and response times as a function of ground truth symbolic distance, now defined in the long axis space (*i*_1_ − *i*_12_). Star indicates significance of the symbolic distance effect; lines show means within the “lower” and “higher” participant partitions. C. Accuracy and RT for each participant (grey dots) in test_long and test_short. Diagonal dashed line is the identity line. Red cross is mean in each condition. D. Choice matrices (upper panels) and RTs (lower panels) separately for the two groups. The lower-performing group exhibit choice matrices which resemble those observed after short axis training, as if they failed to update relational knowledge after boundary training. E. Regression coefficients (*β*s) from fits of idealised test_short and test_long choice matrices (panel A) to human choices (left) and RTs (right). Each dot is a single participant, and is coloured by their accuracy during test_long.

There was substantial variability in performance among participants on test_long, and median accuracy dropped to 79.8%, compared with 88.3% in test_short (Fig. 4C left panel). As average RTs did not differ between test_short (1153 ± 41 ms) and test_long (1166 ± 43 ms) [*t_33_* = 0.49, p = 0.63], this difference was probably not attributable to a decrement in attention between the two conditions (Fig. 4C right panel). Instead, we reasoned that some participants might have failed to fully restructure their knowledge of the transitive series, retaining the belief that the two sets were still independent and treating the relative brispiness of item *i*_6_ < *i*_7_ as an exception. Indeed, participants who performed more poorly (defined by a median split; Fig. 4D right panels) behaved as if they were still in test_short (Fig. 4E, left panel), where as those who performed better generalised the few-shot information about *i*_6_ and *i*_7_ to correctly infer the rank of all other items (Fig. 4D, left panels). Moreover, there was a correlation across the cohort between test_long accuracy and the fit of model test_short behavioral matrices to data obtained from test_long (Fig. 4E, right panel) [choices: *r* = -0.56, p < 0.001; RT: *r* = -0.57, p < 0.001]. We ruled out the possibility that these participants simply failed to learn from the boundary training session, as they still reported the newly trained on object relation - 86 ± 15% of the relevant choices indicated *i*_6_ < *i*_7_ in test_long, compared with 5.8 ± 19 % in test_short for the same cohort (mean ± SD; *t_33_* = 18.0, p < 0.0001; see Fig. 4D. Thus, whilst average participant choices suggested knowledge of a long axis, there was a sizeable cohort that only partially integrated the new relation into their knowledge structure.

Next, we turned to the geometry of neural representations in BOLD during the test_long phase. We considered two hypotheses for how neural representations might adjust following boundary training to permit successful performance on test_long (Fig. 5A). Firstly, under a *hierarchical* coding scheme, the parallel lines observed in test_short (RDM_mag_) might separate along a direction perpendicular to the within-context magnitude axis, so that one dimension codes for a “superordinate” rank given by context (i.e. [*i*_1_ − *i*_6_] > [*i*_7_ − *i*_12_]) and the other for rank within each context (e.g. *i*_2_ > *i*_3_ and *i*_9_ > *i*_10_) [24]. This effectively implements a place-value (or “dimension-value”) representational scheme (akin to numbers in base 6; Fig. 5A, central panel). Note that this hierarchical coding scheme would not require altering the learned neural representation of within-context magnitude, but instead just incorporate additional contextual information, thus predicting increased coefficients for RDM_ctx_. Alternatively, under the *elongation* scheme, objects could be neurally rearranged on a single line stretching from *i*_1_ to *i*_12_ to match our designated ground truth ranking on a single dimension (RDM_mag_long_; Fig. 5A, right panel). We thus constructed a new RDM_mag_long_ that encoded the predictions of this elongation model (Fig. 5C, top panel). Both the *hierarchical* and the *elongation* schemes could potentially allow learning from the boundary training (for items *i*_6_ and *i*_7_) to be rapidly generalised to other items in each context.

**Figure 5.**
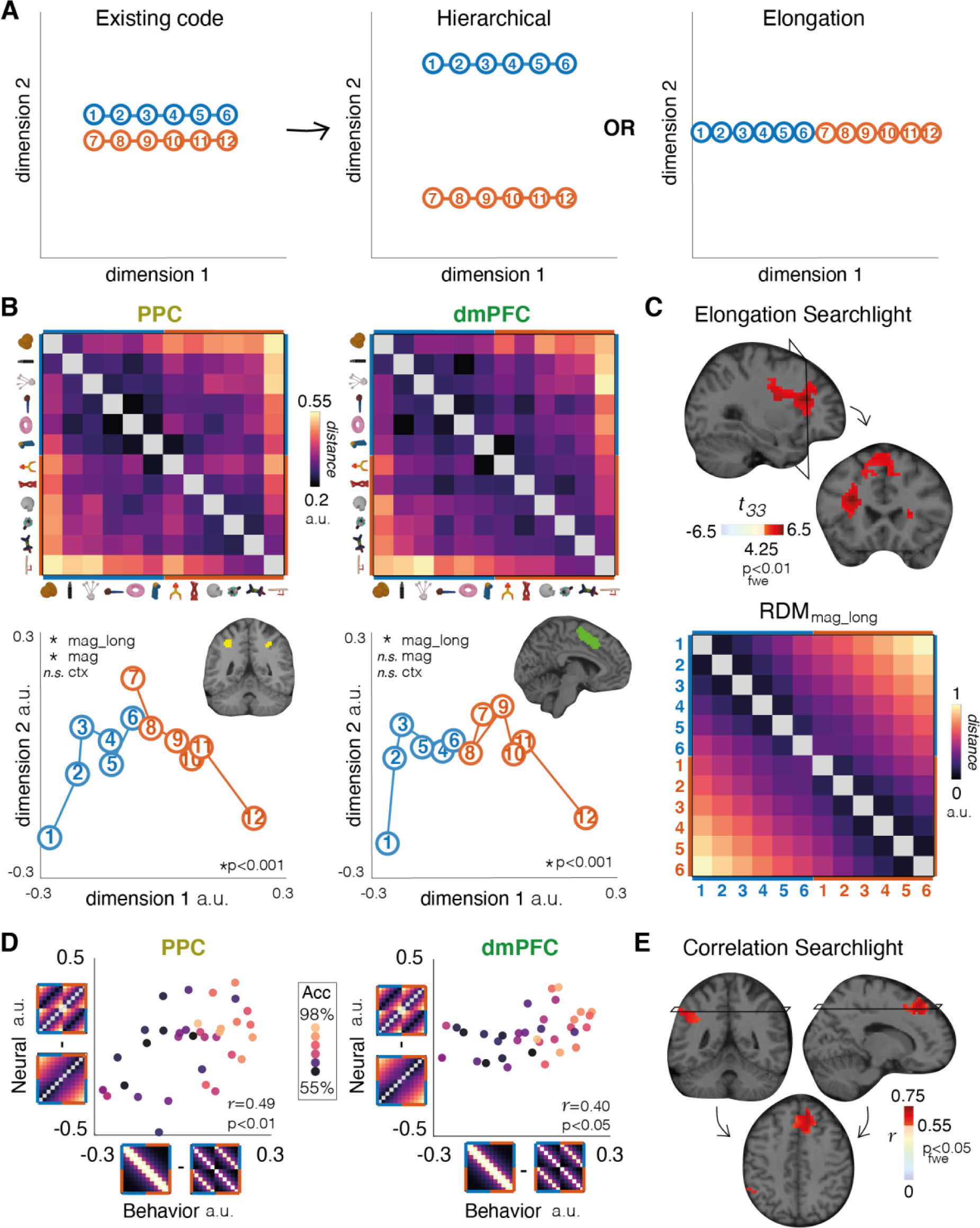
Neural data from test_long. A. Schematic illustration of hypotheses about how the extant neural code (after test_short, right panel) might be transformed after test_long. The hierarchical hypothesis (middle panel) proposes that magnitude and context are represented on factorised (orthogonal) neural axes. Under the elongation hypothesis (right panel), the items are rearranged on a one-dimensional neural manifold (or magnitude line). B. Upper panels: neural RDMs in the PPC and dMPFC after test_long (left panels), and MDS plots (right panels), with ROIs inset. Lower panels: MDS projection of each item in the two contexts (red and blue dots) in test_long for the PPC (right) and dmPFC (left) ROIs. C. Model RDM for magnitude after test_long, and regions correlating with this RDM in a searchlight analysis, rendered onto saggital and coronal slices of a template brain. D. Neural-behavioural integration correlations in PPC and dmPFC ROIs. The x and y axis show relative behavioral model fits (test_long - test_short RT matrices) vs neural fits (elgonation - hierarchical RDMs). The legend displays the relative RDMs (y-axis) and relative RT matrices (x-axis), each rotated into alignment with the axis, from which the neural and behavioural scores were calculated. Each dots is a participant, coloured by their accuracy during test_long E. Voxels showing a significant neural-behavioural correlation

We first compared these schemes empirically by fitting model RDMs to multivoxel pattern data in PPC and dmPFC (Fig. 5B, top row). We compared two regression models, one in which the model RDM was generated under the elongation scheme and one under the hierarchical scheme. Each regession additionally included a predictor coding for RDM_mag_ to accommodate any residual variance due to continued use of a test_short strategy (Fig. S4B). We found that neural data was better fit by the elongation model in both PPC and dmPFC [t_33_ = 4.2, p < 0.001; t_33_ = 5.2, p < 0.001; paired t-test on residual sum of squared error]. Indeed, in both PPC and dmPFC, we observed positive correlations with RDM_mag_long_ [PPC: t_33_ = 4.3, p < 0.001; dmPFC: t_33_ = 6.0, p < 0.001] and when we plotted the neural geometry associated with the 12 items in these regions, it can be seen that they lay on a single (curved) line, consistent with the elongation scheme (Fig. 5B, bottom row; also see Fig. S2). We found no evidence for the hierarchical coding scheme, and in particular no effect of context in our ROIs [PPC: t_33_ = 0.2, dmPFC, t_33_ = 0.6, both p-values > 0.5]. We confirmed the fit of the elongation model in frontal regions using a searchlight approach (Fig. 5C) [peak in left frontal gyrus: -30 23 23, t_33_ = 6.60, p < 0.01 after FWE].

Interestingly, while neural codes in the dmPFC no longer correlated with RDM_mag_ at test_long [t_33_ = 1.9, p = 0.06; t-test on z-scored Pearson correlations with RDM_mag_], the PPC continued to residually code for two overlapping neural lines [*t_33_* = 3.1, p = 0.004; regression with design matrix [RDM_mag_, RDM_mag_long_], t-test on z-scored RDM_mag_ beta weights]. We speculated that this residual coding for the test_short geometry (i.e. parallel lines) may predict the inability of some participants to integrate new knowledge (Fig. 4E). Indeed, we found that participants with a greater tendency to respond as if they were in test_short also displayed a neural geometry more reminiscent of test_short in PPC [*r* = 0.50, *p* = 0.002; pearson correlation between behavioral fits to test_short RT matrix and neural fit to RDM_mag_]. We summarized this relationship by relating the degree of neural elongation (difference in fit for RDM_mag_long_ – RDM_mag_) to the degree of behavioural integration (difference in fit of idealised choice matrices, test_long – test_short) (Fig. 5D left panel) [*r* = 0.49, *p* < 0.01]. We also saw this relationship in dmPFC (Fig. 5D right panel) [*r* = 0.40, *p* < 0.05], but it did not reach threshold in visual cortex [*r* = 0.33, p = 0.06]. Using a searchlight approach within the fronto-parietal network that coded for RDM_mag_ during test_short (see Fig. 3D), we found that this neural-behavioral relationship was expressed most strongly in right Superior Frontal Gyrus (Fig. 5E right; thresholded at FWE p < 0.05, *r* ≥ 0.55, p_uncorr_ < 0.001; also see Fig. S5) [peak correlation: 17 34 54; *r* = 0.71, p_uncorr_ < 0.001; significant at FWE p < 0.01] along with being evident in left parietal cortex (Fig. 5E left) [peak correlation: -48 45 42; *r* = 0.65, p_uncorr_ < 0.001; significant at FWE p < 0.05].

How might knowledge assembly occur on the computational level? Training the neural network with vanilla online stochastic gradient descent [SGD] (as in Fig. 2 and Fig. 3) does not naturally allow the rapid knowledge assembly that is characteristic of human behaviour. In fact, even after pronlonged “boundary” training on item *i*_6_ < *i*_7_, the network learns this comparison as an exception (Fig. 6A), thus failing to generalise the greater (lesser) brispiness to other items in context B (A). In asking how the connectionist modelling framework could be adapted to account for knowledge assembly, one assumption might be that human participants store and mentally replay pairwise associations learned during previous training, intermingling these with instances of boundary training to avoid catastrophic interference [34,35]. However, note that boundary training consisted of just 20 trials comparing *i*_6_ and *i*_7_ with no subsequent rest period that would have allowed time for replay to occur (Fig. 1; also see Fig. S6). Thus, in the final part of our report, we describe an adaptation of SGD that can account for the behaviour and neural coding patterns exhibited by human participants, including the rapid reassembly of knowledge and its expression on a fast-changing neural manifold.

**Figure 6.**
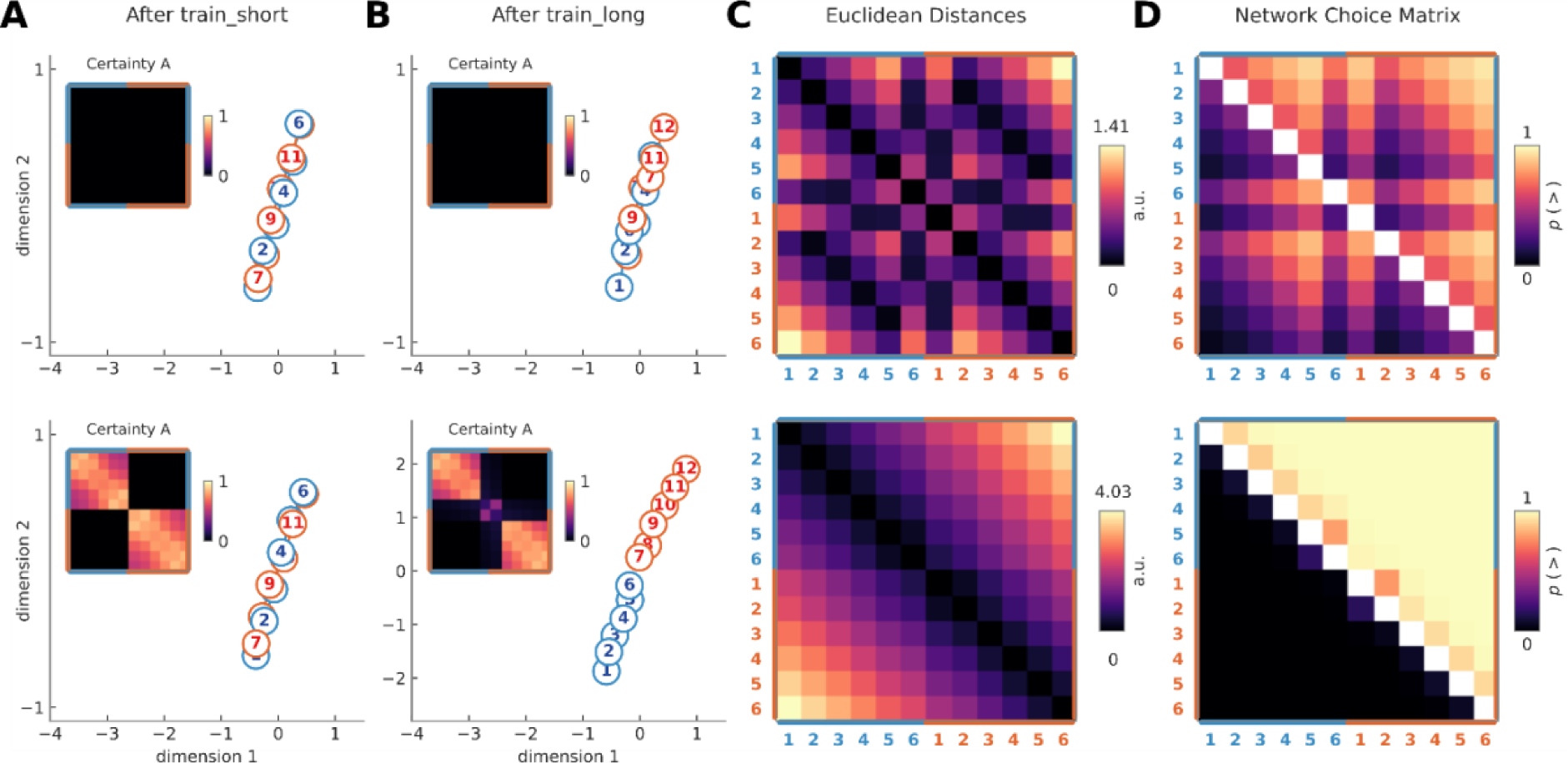
Knowledge assembly in artificial neural network. Fits of neural network to good and poor performers on test_long. Top row: data were generated under the optimal value of *γ* = 2*e*^−3^, which leads to SGD-like training and maximised the fit (mean squared error) to average choice matrices from poor performers (accuracy < median). The two leftmost panels (A and B) show two dimensional MDS of hidden layer representations after train_short (A) and train_long (boundary training) (B) with accompanying certainty matrices (inset). Note the lack of certainty acquistion, suggesting that there is no representational relational among items. After train_short, embeddings for the two contexts (1-6, blue and 1-7 red) lie on two overlapping lines, but these lines are only slightly elongated after train_long (separation between red and blue dots). Bottom row: the same data generated under optimal *γ* = 0.11. This minimal change allows the network to behave like the best human performers. Equivalent matrices for humans are shown in Fig. 4D above.

We reasoned that a simple computational innovation within the neural network could account for the pattern of knowledge assembly observed at test_long. We can think of a neural network as learning to embed inputs on a manifold with maximum potential dimensionality of *d*, equal to the number of hidden units. As we have seen, during training, the network learns to represent stimuli on a manifold with low intrinsic dimensionality (a single neural axis) that represents the transitive series from either context with overlapping embeddings (e.g. Fig. 3A). The assumption we make now is that the network retains a certainty estimate regarding each relation in the embedding space (we call this the *certainty matrix* A). For example, as the relation between items *i*_3_ and *i*_4_ is acquired by the network and the loss consequently decreases, the respective certainty value *A*_3,4_ = *A*_4,3_ increases. With this assumption, new updates can propagate to conserve more certain relations in embedding space, while allowing less certain relations to change. Specifically, gradient updates to the representation of item *k* are mutually applied to all other items *i*_*j*_ ≠ *i*_*k*_, but scaled by certainty *A*_*j*,*k*_ (see Methods). The free parameter *γ* determines the rate at which the certainty matrix is updated as the underlying representations change. This model generalises the case of vanilla SGD which is the special case where *γ* = 0. When training the neural network to solve the transitive inference task, it is possible to recover both successful and less successful knowledge assembly observed in humans by varying *γ* (Fig. 6, Fig. S7B).

We gave the network approximately the same number of train_short and boundary trials as human participants had experienced (960 trials, or 8 cycles, and 20 trials respectively). Over test_short training, the network learned to represent items on two parallel magnitude lines, regardless of the value of gamma (Fig. 6A). Interestingly, performance and training dynamics were indistinguishable across *γ* values after train_short (Fig. S7A). However, the values of the certainty matrix A (inset) for within-context relations still depended on *γ* values. SGD-like training dynamics (*γ* ≈ 0, top row), failed to encode that within-context items are all related to each other in embedding space with certainty, and so the magnitude lines for each context continued to be parallel and overlapping, with the boundary trained items as an exception. By contrast, networks with *γ* ≈ 0.1 (bottom row) learned with high certainty that objects were related within contexts (*i*_1_ − *i*_6_ and *i*_7_ − *i*_12_). As a result, this allowed mutual parameter updates to conserve these relations even with the limited information provided during boundary training (Fig. 6B, lower panel). These updates pushed the contexts in opposing directions, qualitatively consistent with the elongation scheme observed in humans. Interestingly, increasing gamma further causes overly-rapid updates, leading the items of the boundary condition to disconnect from their contexts before the hidden representation were fully elongated (Fig. S7C), suggesting a “sweet spot” for knowledge assembly (Fig. S7B). In fact, we found that we could capture both the high performing human participants and the participants who learned *i*_6_ > *i*_7_ as an exception by varying *γ* (Fig 6D). This exercise revealed good fits for low performers at *both γ* = 2*e*^−3^ and *γ* = 0.87, while only one minima around *γ* = 0.11 fit the participants who correctly assembled the knowledge structures (Fig. S7B; see Methods).

## Discussion

We report behavioural and neural evidence for “knowledge assembly” in human participants. Just 20 boundary training trials were enough for most participants to learn how two sets of related objects were linked. Strikingly, neural representations in multivoxel BOLD patterns rapidly reconfigured into a novel geometry that reflects this knowledge, especially in dorsal stream structures such as PPC and dmPFC. A subset of participants instead considered the boundary relation *i*_6_ > *i*_7_ an exception, and performed more poorly on the subsequent test of transitive inference. while displaying a neural geometry that more consistent with that belief.

The list linking task we use [27] requires participants to make inferences that go beyond the training data – in this case, after boundary training it is parsimonious to assume that because item *i*_6_ > *i*_7_, then item *i*_7_ < *i*_1−6_, and *i*_6_ > *i*_7−12_. How are these inferences made? One possibility is that a dynamic process occurs at the time of inference, proposed by models of transitive inference based on the hippocampus in which updates spread across items via online recurrance. [30]. However, because it takes more cycles to bridge the associative distance between disparate items (e.g. *i*_1_ and *i*_6_), this scheme predicts that more these items would garner longer reaction times and lower accuracy rates– the opposite of the symbolic distance effect we see here.

An alternative is that periods of sleep or quiet resting may allow for replay events, such as those associated with sharp wave ripples in rodents and humans, which might facilitate planning and inference [36], as well as spontaneous reorganisation of mental representations during statistical learning [37]. While we acknowledge that these explanations are not entirely ruled out on the basis of our data, our paradigm allowed very little time for rehearsal or replay – boundary training was few-shot, lasting approximately 2 minutes and comprising just 20 trials in total. Instead, our model proposes that items are earmarked during initial learning in a way that might help future knowledge restructuring, by coding certainty about relations among items (here, a trained transitive ordering). We describe such a mechanism, and show that it can account for our data. Our model is agnostic about how precisely certainty is encoded, but one idea is that in neural systems connections may become tagged in ways that render them less labile. On a conceptual level, this resembles previously proposed solutions to continual learning which freeze synapses to protect existing knowledge from overwriting [38,39]. Thus, notwithstanding a recent interest in replay as a basis for structure memory – including in humans [40-42] – our model has implications for the understanding of other phenomena that involve retrospective reevaluation or representational reorganisation, such as sensory preconditioning [43].

One curiosity of our findings is that unlike for the neural network models, neural manifolds for the transitive series were not straight but inflected around the midpoint (ranks 3/4 in test_short or 6/7 in test_long), forming a horseshoe shape in low-dimensional space. We have previously observed this pattern in geometric analysis of whole-brain scalp EEG signals evoked by transitively ordered images [25] and a recent report has emphasised a similar phenomenon in macaque PPC and medial PFC during discrimination of both faces and dot motion patterns [33]. The reasons for this form to the manifold shape is unclear. One possibility is that the axis coding for choice certainty is driven by the engagement of additional control processes invoked when stimuli evoke conflicting responses, and that these proceses are currently missing from our neural network model [44]. Another is that the horseshoe shape allows neighbouring items to be linearly discriminated, by the judicious application of hyperplanes with gradually varying angle with respect to the neural manifold. Resolving this issue is likely to be an important goal for future studies.

In sum, we observed rapid recorganization of neural codes for object relations in dorsal stream structures, including the PPC and dmPFC. This is consistent with a longstanding view that dorsal structures, and especially the parietal cortex, encode an abstract representation of magnitude or a mental “number line” [7,45,46]. Recently, many studies have emphasised instead that the medial temporal lobe, and especially the hippocampus and entorhinal cortex, may be important for learning about the structure of the world [3,6,22,47]. One important difference between our work and many studies reporting MTL structures is that our study involved an active decision task (infer brispiness) whereas previous studies have used passive viewing or implicit tasks to measure neural structure learning. It may be that dorsal stream structures encode structure most keenly when relevant for an ongoing task. We do not doubt that both regions are important for coding relational knowledge, but their precise contributions remain to be defined.

## Online Methods

### Participants

Thirty-seven healthy adult participants were recruited for this study. One was excluded for failure to reach performance threshold (see above), and two more for practical reasons (failure to attend scanning session; discomfort in scanner leading to early termination of the experiment). This left n = 34 total (19 males, mean age: 23.3 ± 3.4 years). Participants reported no history of psychiatric or neurological disorders, and gave informed consent prior to scanning. The study was approved by the ethics committee of the University of Granada. Participants’ base compensation was 35 Euros, plus a performance-based bonus for an average payment of 40.93 ± 2.57 Euros. Participants were given a voluntary anonymous debrief concerning their insight into the test_long session, which we display in Supplementary Table 1.

### Stimulus and task

Stimuli were novel objects drawn from the NOUN database [28]. Out of the 60 possible images in this database, objects that were rated as most similar to the others (e.g. a similarity rating within 1 standard deviation of the maximum) and objects that were rated as most familiar (e.g. less than 50% for inverse familiarity score) were excluded, resulting in 41 possible objects. For each participant, 12 of these 41 objects were randomly selected for use throughout the study. Selected objects were arbitrarily assigned a rank from 1-12, with ranks 1-6 belonging to context A and 7-12 to context B. We denote these *i*_1_ − *i*_12_ in the text.

Before entering the scanner, participants performed a computerised training phase which we call train_short. This training phase consisted of between 3 and 10 blocks of 120 trials. On each block, objects were sampled from a single context for 60 trials (A or B) and then the alternate context for another 60 trials. Each trial began with the presentation of two objects drawn from adjacent ranks withinin a single context (e.g. *i*_3_ and *i*_4_ or *i*_8_ and *i*_9_), which were shown either side of a central fixation point. Above the point the words “more brispy?” or “less brispy?” appeared in Spanish (i.e. “mas brispo?” or “menos brispo?”). These objects remained on screen for 5000 ms or until response, whichever was shorter. Participants were instructed to select the corresponding object (i.e. that which was more or less brispy) using either the “F” (left object) or “J” (right object) keys. Once a response was recorded, a red or green box would appear around the selected object to indicate whether it was the correct selection, and this response feedback box persisted for 475 ms. If participants did not respond within 5000 ms, the trial was considered incorrect and was not repeated. After feedback there was a blank screen for a variable delay of up to 50 ms before the next trial. Critically, participants were only trained to compare 5 object pairs from each context, e.g. *i*_1_ − *i*_2_, *i*_2_ − *i*_3_, *i*_3_ − *i*_4_, *i*_4_ − *i*_5_, and *i*_5_ − *i*_6_, from context A and *i*_7_ − *i*_8_, *i*_8_ − *i*_9_, *i*_9_ − *i*_10_, *i*_10_ − *i*_11_, and *i*_11_ − *i*_12_, from context B.

Whether participants were asked to select the more or less brisp object, and hemifield presentation of the objects, were randomised on each trial. Additionally, the trial-order of each object pair was randomly shuffled. Participants performed this task for at least 3 blocks, and until they reached a criterion of 90% correct on both contexts and correct responses for the final 12 comparisons of each context. One participant was excluded for failing to reach this criterion after 10 blocks (see Fig. 1C).

The format of the test phases test_short and test_long was identical (but different to train_short). Test phases occurred in the scanner, and consisted of 288 trials in which lone objects were presented in a random sequence, with the constraint that each combination of 12 (current trial) x 12 (previous trial) ranked objects occurred exactly once in the first half (144 trials) and once in the second half (144 trials) of the test phase. Each object was presented centrally for 750 ms, after which participants had 2000 ms to respond whether it was more or less brispy than the previous object. The words “more” and “less” appeared randomly on the left and right of the screen, and the mapping from more/less to left/right buttons (held in either hand) switched midway through the test phase. After the response, there was a pseudorandomly jittered interval of 1500-5500 ms, and the fixation dot turned blue if a response was recorded within this deadline, while a red letter X appeared if the response was missed. Critically, participants did not receive trial-wise feedback and were instead were rewarded bonus points at the end of each block. These bonus points were proportional to their accuracy on that block and were translated into additional monetary reward at the end of the experiment.

The boundary training phase occurred between test_short and test_long. It was similar to train_short except that it lasted just 20 trials, and the only items presented were objects ranked (arbitrarily) as 6 and 7. These could occur on either side of the screen, with “more” or “less” randomised over trials as in train_short. Participants viewed the objects for 3000 ms after which a feedback screen stayed up for 1500 ms, be it a green or red bounding box, or a red X at fixation if no response was recorded. There was then a variable intertrial interval from 1400-5000 ms before the next trial.

Finally, after completing test_long, participants remained in the scanner and performed a number localiser task, which was identical to test_short / test_long with the exception that objects were replaced with Arabic digits 1-12 and participants responded “more” or “less” according to whether each number was greater or less than the previous.

### Idealised Behavioral Matrices

We analyses behavioural data by plotting accuracies and RTs for each combination of 12 objects shown at test, and/or as a function of symbolic distance (i.e. the distance in rank between the current and previous item). We constructed idealised reaction time and choice matrices (Fig. S4A) under the assumption that choices were noiseless triangular matrices and that RTs depended linearly on symbolic distance. Specifically, our RT matrices were constructed by 1⁄(1 + *dist*(*v*′, *v*)), where *v* = [1: 6 1: 6] for test_short and [1: 12] for test_long, and *v*’ it’s transpose. Our accuracy matrices were created by setting the upper triangle of each quadrant (in test_short) or the entire matrix (in test_long) to 1 and the lower triangle to zero. Note that as participants did not compare objects to themselves, diagonal elements of our design matrices were excluded from analyses.

### fMRI data acquisition

MRI data were acquired on a 3T Siemens scanner. T1 weighted structural images were recorded directly prior to the task using an MPRAGE sequence: 1x1x1 mm^3^ voxel resolution, 176x256x256 grid, TR = 2530 ms, TE = 2.36 ms, TI = 1100ms. Each fMRI image contained 72 axial echo-planar images (EPI) acquired at a multiband acceleration factor of 4 in interleaved sequence. Voxel resolution was 2 mm^3^ isotropic, slice spacing of 1.6 mm, TR = 1355 ms, flip angle = 8, and TE of 32.4 ms. 560 EPI images were recorded for the localiser and 1220 EPI images for each of the experimental sessions. This resulted in 3000 EPI images per participant with a scanning time of about 100 min. Scans were realigned to the mean scan within each session. The anatomical scan was co-registered to the mean of all functional images. Anatomical scans were normalized to the standard MNI152 template brain. The functional EPI images were then normalized and smoothed with a full width half maximum Gaussian kernel of 8mm. Images were then downsampled by reslicing to 3 x 3 x 3 voxel mm^3^ voxel resolution before performing analyses.

### fMRI: Data analysis

Scanning sessions were concatenated and constants included in the GLMs to identify runs and account for differences in mean activation and scanner drift. All GLMs used delta functions convolved with the canonical haemodynamic response function (HRF) and time-locked to trial events. We also included the 6 head motion parameters derived from pre-processing as nuisance regressors [translation in x, y, z; yaw, pitch, roll]. Automatic orthogonalization was switched off. Data were analyzed with SPM12 and in-house scripts. All contrasts were constructed as simple t-contrasts with first-level t-maps as input. Unless otherwise noted, we only report clusters that fell below an FWE-corrected p value of 0.01 (as in [48]) with a setting of cluster extent to 10 voxels or more and a voxel-wise uncorrected threshold of p < 0.001. Data were visualized using the XjView toolbox (http://www.alivelearn. net/xjview).

We fit our data with 4 different general linear models (GLMs). The first GLM was used to define ROIs from the number localiser. The design matrix for GLM1 included parametric modulators time-locked to stimulus onset for each number, as well as 6 nuisance motion regressors. We considered clusters of voxels that passed a threshold of FWE p < 0.01 in response to the stimulus regressor. Althought several regions passed this threshold, we focussed on ROIs in dmPFC and PPC, chosen on the basis of previously stated predictions (we show searchlight results in addition to ROI analyses, which seem to justify this choice) [7]. The second and third GLMs were used to estimate neural patterns associated with each object within test_short and test_long. The design matrix for these models each included 12 regressors, one for each of the objects locked to stimulus onset, as well as 6 additional nuisance regressors for head motion. In one case (GLM2) we estimated this regression separately for each block of 72 trials (n = 4). This allowed us conduct analyses that required between-run crossvalidation (e.g. SVM analysis). In the other case (GLM3) we modelled all trials within test_short or test_long. This latter GLM was used for calculating RDMs in ROI and searchlight analyses of fMRI data. In a fourth GLM, we additionally included either 11 (test_short) or 22 (test_long) regressors coding for the distance from the current to previous image. Fits from GLM4 were used to generate data for multidimensional scaling visualization aids.

### Representational Similarity Analysis

BOLD RDMs were constructed by taking the correlation distance between voxel patterns elicited by each of the objects in test_short and test_long, yielding a 12 x 12 RDM. For searchlight analyses, we used a radius of 12mm. For each searchlight sphere or ROI, we computed the neural DMs from mthe condition-by-voxel matrix of estimated neural responses using Pearson corelation distance between pairs of conditions.

These were compared to model RDMs which were created from linear distances between item ranks within context (1-6 and 7-12; RDM_mag_), distances between ranks across contexts (1-12; RDM_mag_long_) and between contexts themselves (i.e. 0 within context, 1 between context). All model RDMs were standardised and comparisons to neural data were conducted with tests of correlation (pearsons *r*), or regression. The additional RDM reported here (RDM_NN_) was obtained by taking the Euclidean distance between the 12 hidden-layer activations elicited by probing the network with one-hot inputs corresponding to *i*_1_ − *i*_12_). Z-scored neural RDMs were regressed, or correlated, with z-scored mmodel RDMs. All statistics reported for RSA analyses were obtained by evaluating RDMs at the single subject level and conducting group-level (random effects) inference on the resulting coefficients, using FWE correction where appropriate.

To visualise neural state spaces, we used multidimensional scaling with metric stress (equivalent to plotting the first principal components of the data) in two dimensions, using GLM4 (see above).

### Support Vector Machine Decoding

All classification analyses utilised a multiclass support vector machine model. GLM2 utilised 4 beta values for each object (one for each run), and so we trained binary SVM classifiers on data from objects in context 1, and tested the model on objects in context 2. The classifier used a “one versus one” coding design, meaning that each learner *l* was trained on observations in 2 classes, treating one as the positive class and the other as the negative class and ignoring the rest. To exhaust all combinations of class-pair assignments, we fit 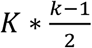 binary SVM models, where *k* are the unique classes (ranks 1-6 here). Specifically, let *M* be the coding design matrix with elements *m*_*kl*_, and *s*_*l*_ be the predicted classification score for the positive class of learner *l* (without loss of generality). The algorithm assigns a new observation (from context 2) to the class ^*k̂*^ that minimizes the aggregate loss for the *L* binary learners.

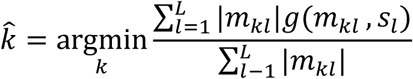

Each of these binary learners used a linear kernel function.

### Brain Behavior Correlations

We performed a correlation analysis to quantify the extent to which the elongation of neural representations predicted integrated behavioral responses. We analyzed human choice patterns by computing idealsed choice matrices (described above) and inputting both the test_short and test_long patterns into a competitive regression model. The degree of behavioral integration was defined as the the relative fit of each of these matrices. Similarly, we constructed neural model RDMs describing the ground truth symbolic distance between each pair of item in test_short and test_long (see above). We then defined the degree of neural elongation as the relative fit to each of these RDMs in a competitive regression model. We then tested at the group level the extent to which the degree of neural elongation predicted the degree of behavioral integration using pearsons correlation.

### Neural Network Simulations

We implemented a two-layer feedforward neural network with coupled input weights to study the computational underpinnings of the task, its solutions and possible failure modes. All simulations were run for *n* = 20 random seeds and plots show averages across seeds. The network received two one-hot vectors, coding respectively for the object on the left (*x*_*a*_) and right side (*x*_*b*_) of the screen (a one-hot vector for object *i* has zeros everywhere except at the *i*-th position which is equal to one). The one-hot vectors were then propagated forward by the coupled weights *W_1_* and -*W_1_* respectively, followed by a rectified linear unit (*ReLU*) to create the hidden layer representation of 20 neurons. Finally, the hidden layer representation was projected onto a single output value by the readout weights *w2*:

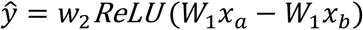

By coupling the input weights we ensured that the hidden layer representation of each object was independent from the position on the screen. Note that due to objects being represented as one-hot vectors, hidden representations of objects are independent from each other, i.e. the *i*-th column of the weight matrix *W*_1_. Further, due to the symmetry in the first-layer weights, if the two one-hot vectors encode the same object the network’s output is zero.

The network was optimised using stochastic online gradient descent

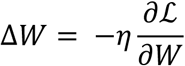

on single pairs of objects, i.e. a batch size of 1, with learning rate η = 0.05 on the mean squared error between the network’s output *ŷ* and the target values *y* = 1 for *i*_*a*_ > *i*_*b*_ and otherwise *y* = −1:

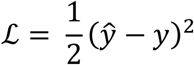

Since inputs to the network were two one-hot vectors, in each training step only two columns of the first layer weight matrix *W*_1_ were updated, we denote these two column vectors by Δ*w*_1*a*_ and Δ*w*_1*b*_.

Synaptic weights were initialised from a zero-centered Gaussian distribution with standard deviation 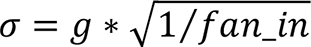 where *g* = 0.025 and *g* = 1 in hidden and readout layers respectively. The hidden layer weights were initalised to small values to encourage a low-dimensional (“rich”) solution [32]. We employed a training procedure very similar to that used for human subjects. Networks were first trained for 8 cycles, where each cycle was compmrised of 120 trials (60 per context) - leading to a total of 960 steps of gradient descent training. Subsequently, we performed 20 training steps on the two objects of the boundary condition.

### Learning relational certainty

In order to recover the rapid *knowledge assembly* observed in humans, we adapted vanilla SGD, by applying mutual updates on synaptic weights *W*_1_ based on the pairwise certainty that two object representations bear an accurate relation to one another in embedding space, and in addition, correcting for potential drift in the readout weights *w*_2_. For a given trial *t* with inputs *x*_*a*,*t*_ and *x*_*b*,*t*_ we compute the certainty value as a sigmoidal function of the loss incurred ℒ_*t*_, with the slope α and the bias *β* of the sigmoid as potentially free parameters (here we set them to *α* = 1000 and *β* = 0.01)

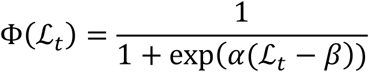

Pairwise certainty values were then stored in the certainty matrix A as an exponential moving average

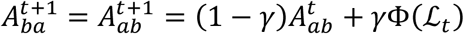

where the free parameter *γ* determined how quickly old values were discounted.

In addition, to infer certainty values for pairs of items that were not presented to the network, a fraction of the certainty values for item *i*_*b*_ were added to the certainty values of *i*_*a*_.

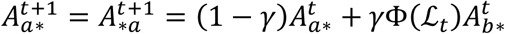

and vice versa. Note that the certainty matrix is symmetric and therefore rows *A*_*a*∗_ are identical to columns *A*_∗*a*_. These row-wise updates followed the heuristic: If item *a* is correctly related in embedding space to item *c* and item *a* is correctly related to item *b*, then infer that item *b* is also correctly related to item *c*. Synaptic weights were then mutually updated by outer products:

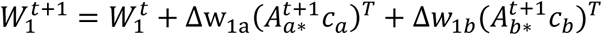

where *A*_*a*∗_ denotes the *a*^th^ column of the certainty matrix. Note that *c*_*a*_ and *c*_*b*_ are vectors of scaling factors to correct for drift in the readout weights as follows:

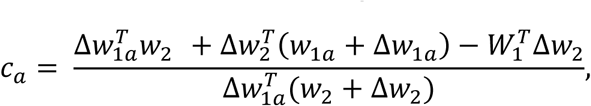

Ins order to perform a SGD step on the items currently presented to the network (i.e. the a-th and b-th column of *W*_1_) the *a*-th and *b*-th entry of *A*_*a*∗_, *c*_*a*_ and *A*_*b*∗_, *c*_*b*_ respectively are set to 1.

### Fitting Human Choice Matrices

To fit human choice matrices we applied a sigmoid function to the linear neural network output

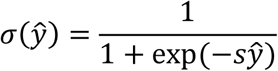

We fit different parameterisations separately to choice matrices for high and low performers (defined by a median split, as in Fig. 4). For each, we performed a grid search on combinations of *γ* in range 0 to 1 and s in range 0.01 and 100 (both in log10 units) and mapped the resulting deviation between predicted and observed choice matrices for that participant group (Fig. S7B). Because neural network models are stochastic, we repeated the simulations for 20 random initial seeds and averaged the resulting deviance for fitting. Low performers were fit well with *s* ≈ 1 and had a U-shaped relationship with accuracy for varying *γ*, leading to two local minima, such that values that were close to zero (vanilla SGD) and close to one both resulted in a failure to stitch information appropriately (Fig. S7B). For low *γ*, tha algorithm fails to acquire certainty and thus does not perform mutual updates, behaving like vanilla SGD. Similarly, for large *γ*, the certainty matrix is rapidly udated, such that the boundary items form a high certainty cluster separate from the rest of the items (Fig. S7C). In this case, after few mutual update steps that partly disentangle the representations, the certainty matrix rapidly approaches zero for non-boundary items, again leading to SGD-like updates on boundary items only. Behaviourly, these two failure modes can be interpreted as either failing to relate items in the two conditions during after boundary training for low *γ* or by relating the items of the boundary condition as independent group during train_short for large *γ*. On the other hand, high performers had a single minimum for *s* > 2 and *γ* ≈ 0.1 (Fig. S7B).

### Supplementary Materials

**Supplementary Figure 1.**
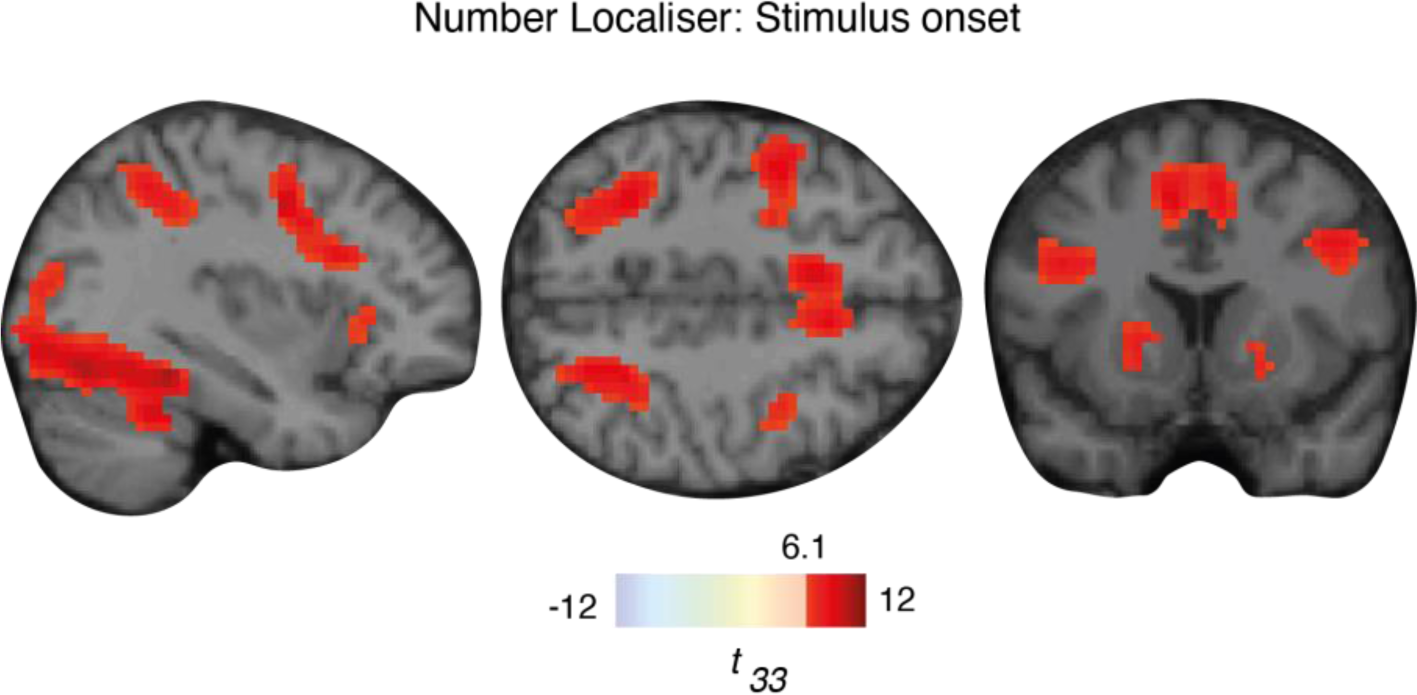
Univariate activation in Number Localiser. Three views (sagittal, axial, coronal) of maps of significant BOLD response to stimulus onset (Arabic digits 1-6) during the number localiser task. This independent task was used to define regions of interest (ROIs) for our main analyses (Fig. 3E). Overlays indicate voxels showing a main effect of stimulation in the localiser task at the group level (GLM1; see Methods). We use a statistical threshold determined using a familywise error (FWE) threshold of p < 0.01, corresponding to t_33_ > 6.1.

**Supplementary Figure 2.**
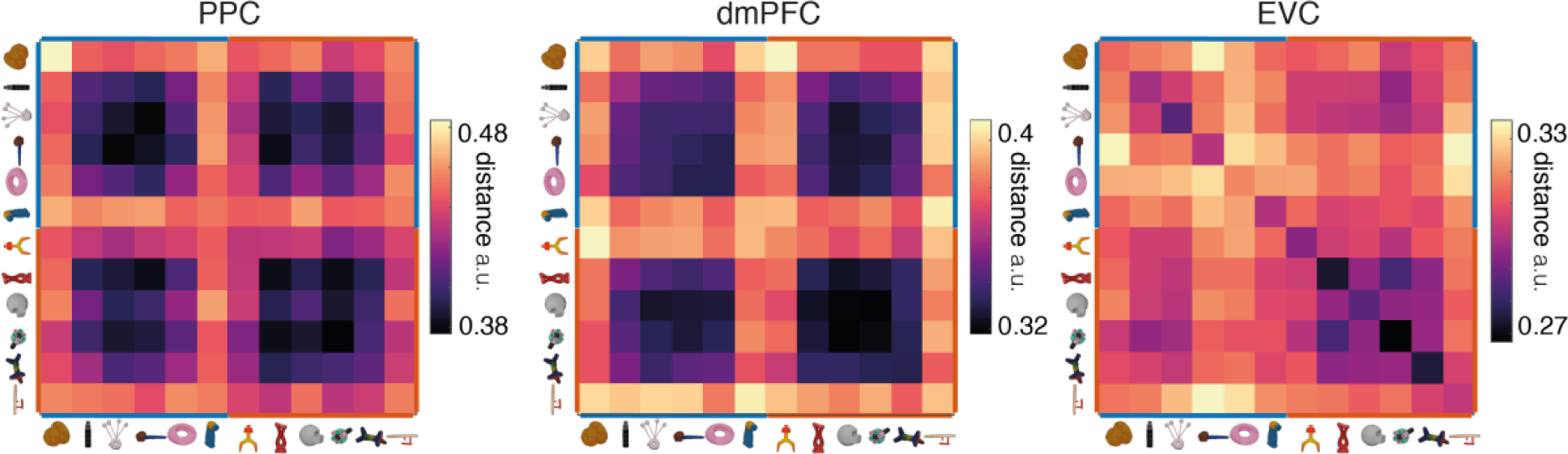
Cross-validated Neural RDMs. These RDMs were computed by taking neural pattern (correlation) distances between each of the 12 objects in each run and every other (nonidentical) run. Thus, the diagonals are not necessarily zero in this analysis. This also allows us to show exemplar discriminability across sessions, i.e. the similarity between each object and itself across runs, which is highest in extrastriate visual cortex (EVC; rightmost plot). As there are 4 sessions, each of these plots is the average of all 6 possible cross-validated distances 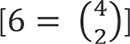; purple values indicate lower distances, while yellow values indicate higher ones. As we counterbalanced responses within participants (see Methods), these plots average over experimental sessions with opposing response mappings.

**Supplementary Figure 3.**
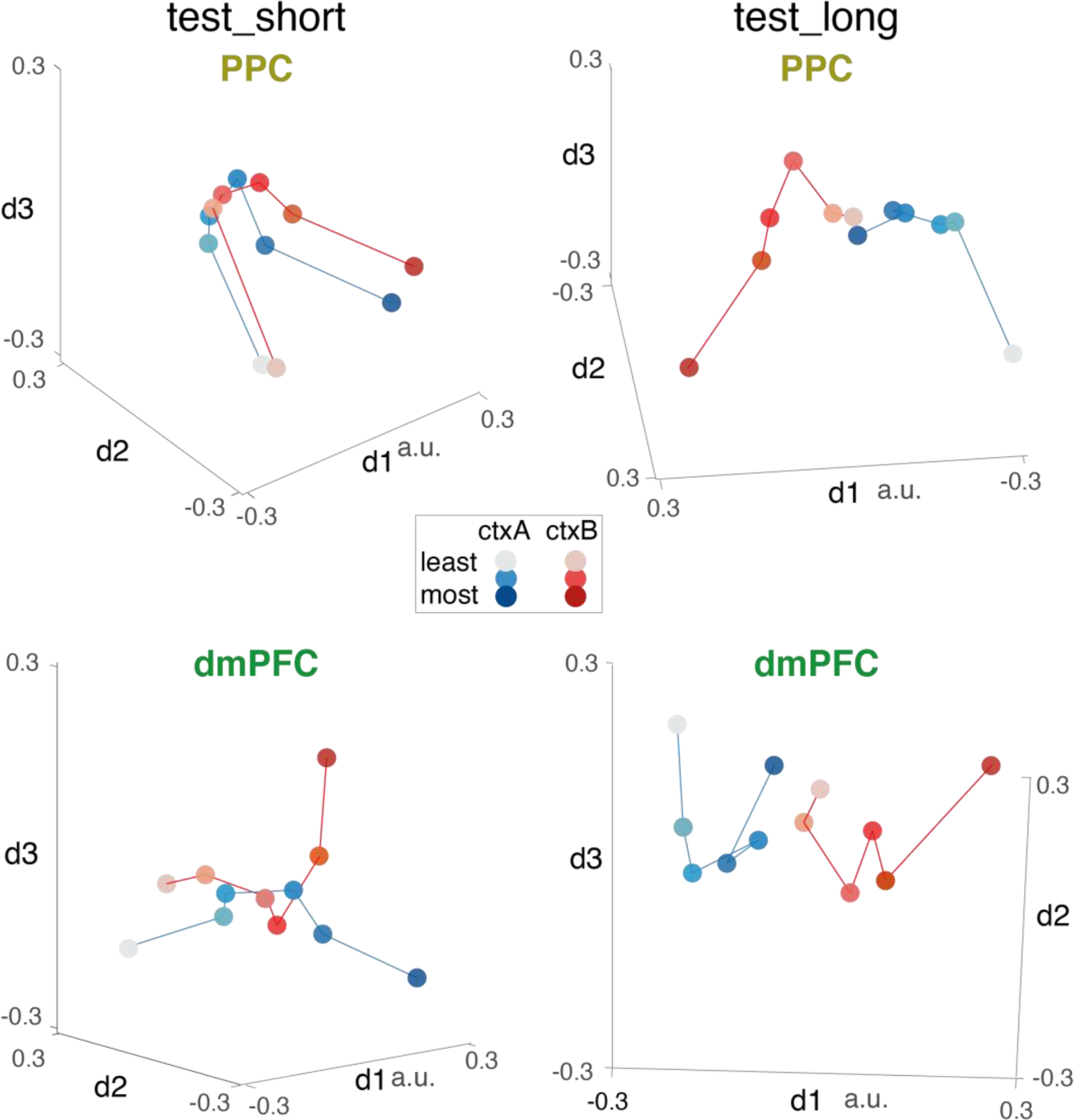
Multidimensional scaling plots in ROIs. Supplemental multidimensional scaling views of BOLD representations during test_short (left panels) and test_long (right panels) from the PPC and dmPFC regions of interest. Each axis shows one of the three dimensions identified by MDS. Axis rotation is different in each plot and chosen for illustrative purposes. Each dot is a stimulus, shaded by its rank from “most brispy” (darker colors) to “least brispy” lighter colours. Items from context A (ctxA) are shown in blue and context B (ctxB) in red. Distances between circles approximate similarities in the RDM (in low rank). These plots correspond to the 2D MDS plots reported in the main text (Fig. 3C and Fig. 5B); they are provided as visualization aids – all statistical analyses were performed on the full-rank representational dissimilarity matrices.

**Supplementary Figure 4.**
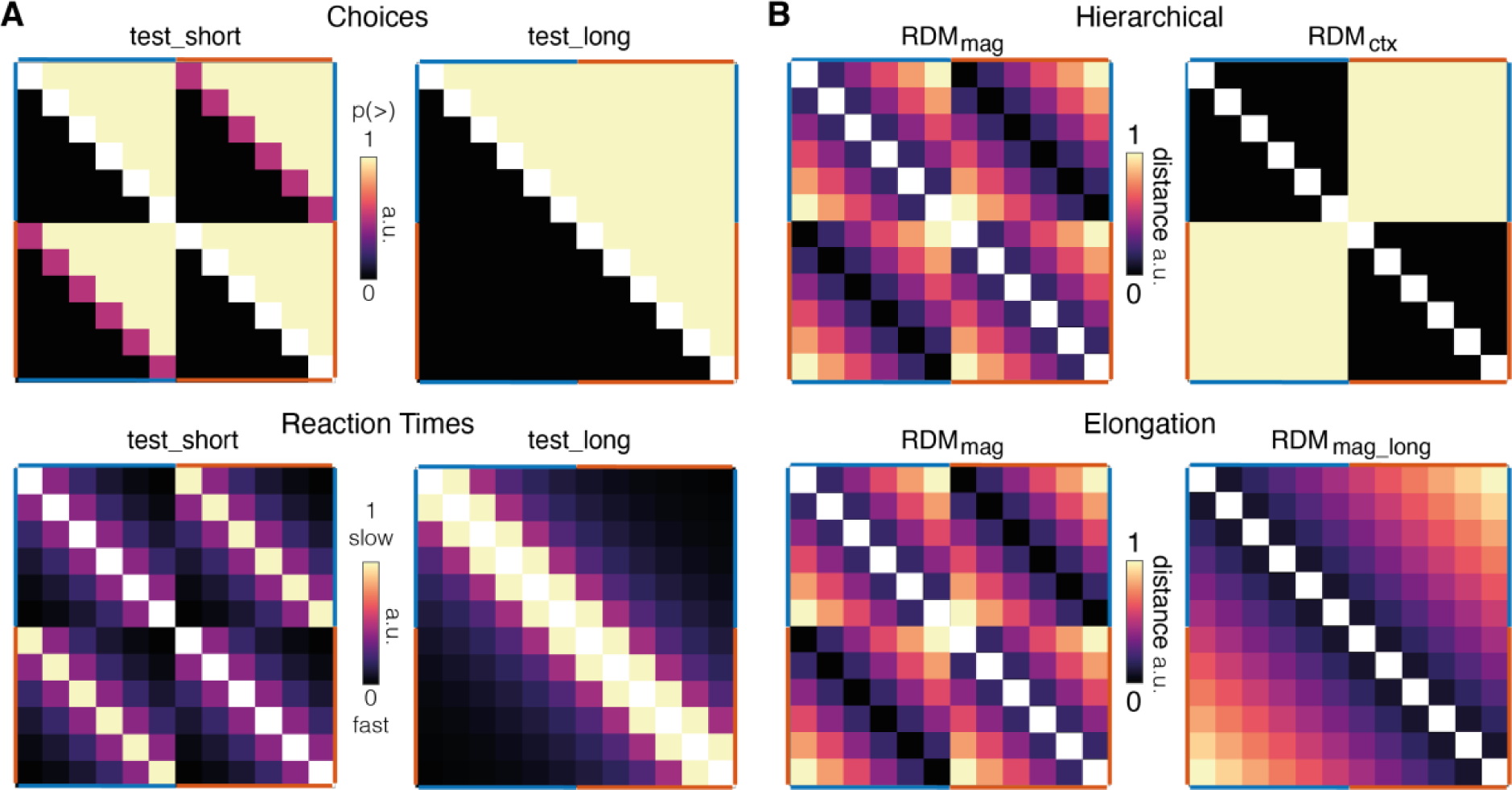
Idealised behavioral matrices and Neural RDMs. A. Idealised behavioral matrices for choice (top row) and reaction times (RT; bottom row) used for test_short (left column) and test_long (right column). The choice matrix has values of 1 for each ground truth “more” response, 0 for each ground truth “less” response, and 0.5 for each ambiguous response. The RT matrices are scaled linearly with symbolic distance, ranging from 0 (fastest RT for comparisons with symbolic distance = 5) to 1 (slowest RT for comparisons of symbolic distance = 0). These idealised matrices were used to quantify Human participant behavioral choice and reaction time patterns (Fig. 4), and so diagonal axes were set to empty values (indicated with white cells). B. Test_long neural hypotheses: boundary training could have altered neural codes to accommodate new information in several ways. Here, we plot the idealised predictor RDMs utilized in regression models testing the Hierarchical (top) vs Elongation (bottom) candidate theories of knowledge assembly hypotheses reported in the main text (Fig. 5).

**Supplementary Figure 5.**
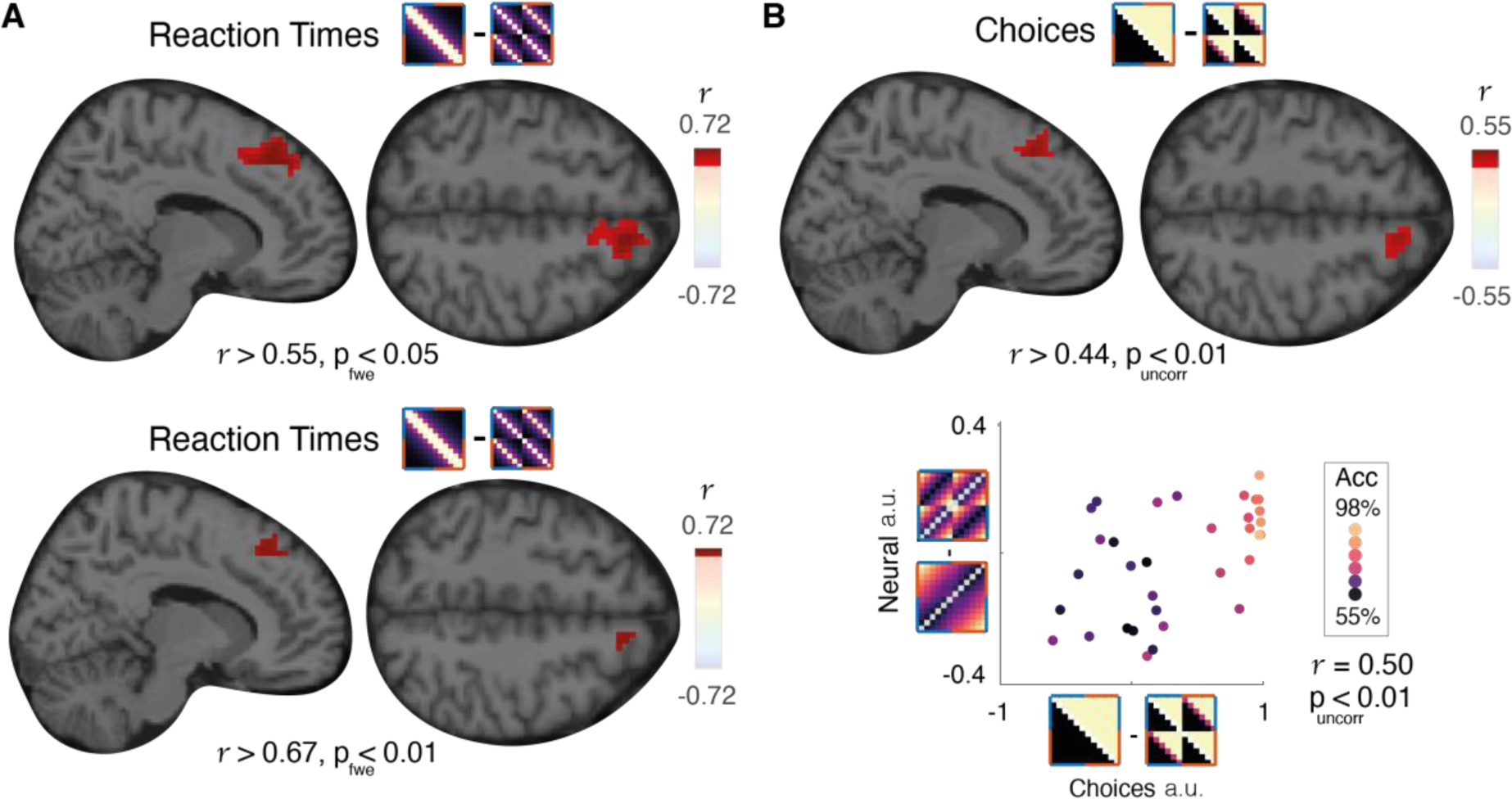
Brain-behaviour correlations. A. Top row: Maps of the brain-behaviour correlation across participants between the difference in fit between model RDMs (RDM_mag_long_ - RDM_mag_) to neural data and the difference in fit between idealised choice matrices (test_long - test_short) thresholded at p_fwe_<0.05 – the same threshold displayed in Fig. 5E. Bottom row: Similar to A, but now thresholded at p_fwe_<0.05. Note that all brain images were estimated with a searchlight approach and display MNI coordinates [11 33 51]. B. Top panels: Brain-behaviour correlations defined by choice matrices thresholded at p_uncorr_<0.01. This cluster was no longer significant after correction at p_fwe_<0.05. Note that all brain images were estimated with a searchlight approach and display MNI coordinates [11 33 51]. Bottom panel: We plot the neural-behavioral correlation within the cluster shown in the top row. The legend displays the relative RDMs (y-axis) and relative choice matrices (x-axis), each rotated into alignment with the axis, from which the neural and behavioural scores were calculated. Each dot is a participant with a colour that denotes its accuracy. The main text (Fig. 5E) shows the equivalent for idealised RT matrices (also see Fig. S4 above).

**Supplementary Figure 6.**
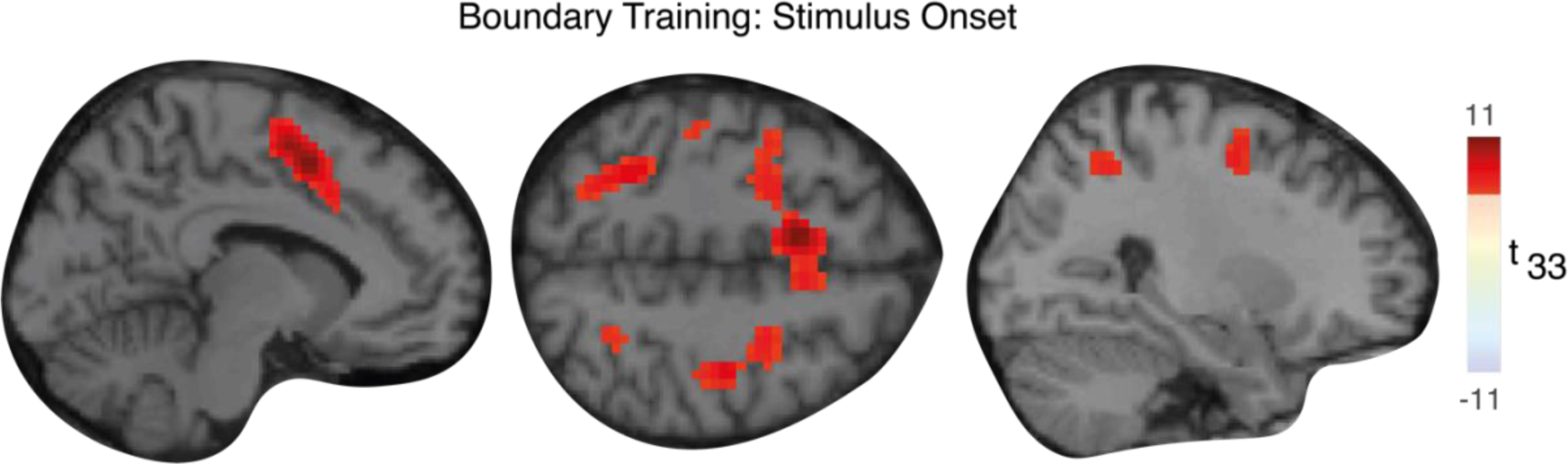
Univariate activation in response to stimulus onset during boundary training. Map of the regions that showed significant activation in response to the onset of the two boundary items during the 20 learning trials (train_long). Brain maps are thresholded at p_fwe_ < 0.05, corresponding to t_33_ > 5.67. We observed peaks in left medial frontal [MNI coordinates: -9 6 51, t_33_ = 11.0] and right frontal cortex [MNI coordinates: 42 9 30 t_33_ = 9.9, p_uncorr_ < 0.0001] but a lack of hippocampal activation (t_33_ < 2.5 over bilateral hippocampus and bilateral parahippocampal regions). Additionally, hippocampus did not appear to code for context in test_long (*r* = 0.41), similar to results observed in PPC (*r* =0.22) and dmPFC (*r* =0.6, all p>0.5)

**Supplementary Figure 7.**
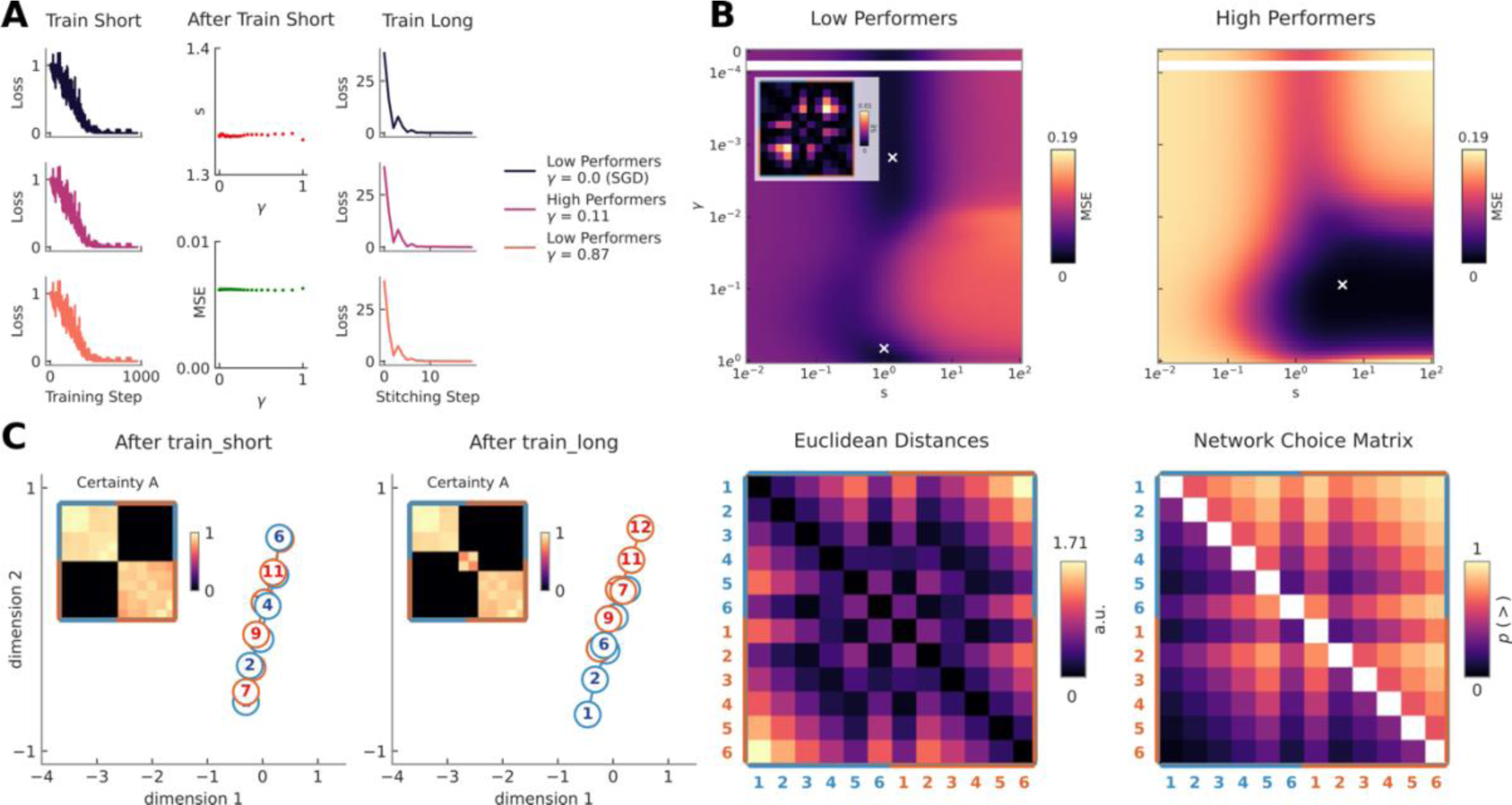
Fitting low and high performers. A. Left column: Average training loss during train_short averaged across n=20 randomly initialized seeds of the neural network for different values of gamma (see legend, right) are indistinguishable. Middle column: When fitting the average human choice matrix after train_short, the optimal s and resulting mean squared error (MSE) are stable across values of gamma (central panel), further indicating that behaviour is indistinguishable after initial training across low and high performers. Right column: Average training loss during train_long across n=20 seeds is indistinguishable between different values of gamma (see legend, right). B. Mean squared error of behavioural fits for low (left panel) and high performers (right panel) across gamma and s values in log units. Minima are indicated by white x’s. Fits to average choice matrix of low performers reveal two minima, both around s=1 and either at low or high gamma (s=1, gamma=0.002; s=1, gamma=0.87). We also display the squared difference between the fitted behavioural matrices for the two local minima (inset). The average choice matrix of high performers is fit well for low gamma and s>2 (s= 4.87, gamma=0.11). C. Here we show the same plots as in Fig. 6 but for the second local minima of low performer’s fit (s=1, gamma=0.8). We plot the two-dimensional MDS of hidden layer representations after train_short (first panel) and train_long (second panel) with accompanying certainty matrices (insets). We show the Euclidean distances for all combinations of items across both contexts (third panel), as well as the best fit neural network choice matrix to low performers’ behavior (fourth panel)

### Normalisation

Since an elongated brispiness axis was observed during both test_short and test_long in ROIs from our number localiser (see Methods), we next asked about the relationship between these representations *across tasks*. To do this, we fit a GLM to data jointly across all three experiment types. As with those reported in the main text, this GLM used delta functions convolved with the canonical haemodynamic response function (HRF) and time-locked to trial events. The design matrix for this model included 30 regressors, one for each of the Arabic digits or objects locked to stimulus onset, as well as 6 additional nuisance regressors for head motion. We then constructed RDMs as in the main text to assess the similarity of evoked neural patterns, this time focusing on how patterns were related *across* the experiments (Fig. S8A).

We hypothesized that neural populations could code for either ground truth magnitude, in which case we would expect the test_long axis to be twice as long as in test_short and the number localiser (RDMnone, Fig. S8B), or for relative magnitude, meaning the axes in all three experiments should be compressed to the same length (RDM_sub,div_, Fig. S8B). Specifically, for RDMnone, distances were computed between the following arrays: [1: 6] in the number localiser and in test_short and [1: 12] in test_long. We also constructed an idealized matrix under a subtractive normalization scheme, RDMsub, that computed distances between arrays for the number localiser and test short, constructed as [1: 6] − mean([1: 6]), as well as test_long: [1: 12] − mean([1: 12]). Finally, we also derived an idealized matrix under a subtractive and divisive scheme, RDM_sub,div_ in which we computed distances between the following arrays 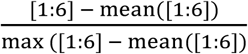 (used for the number localizer and test_short) and 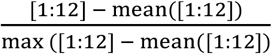 comparison. (test_long). All model and neural RDMs were z-score normalized before

We found evidence for this normalised coding scheme in both PPC and dmPFC (Fig. S8B; one-sided t-tests on Pearson correlations: RDM_sub,div_ t_33_ = 3.86, 6.29, p’s<0.001; RDMnone t_33_ = -3.07, -0.5, p’s>0.95; RDMsub t_33_ = -1.8, 0.15, p’s > 0.6; PPC and dmPFC respectively). This type of normalisation has previously been observed in other settings [24], and its potential role in knowledge assembly is an intriguing point for further investigation.

“In the second learning session, you learned that there were two groups of objects were connected. Specifically, the most brisp object of one group was no less than the least brisp object of the other group. Did you notice that? Was it clear that all the objects were part of one order?

Even if not, tell us here if and how you thought the objects were related after the second learning session.”

**Supplementary Figure 8.**
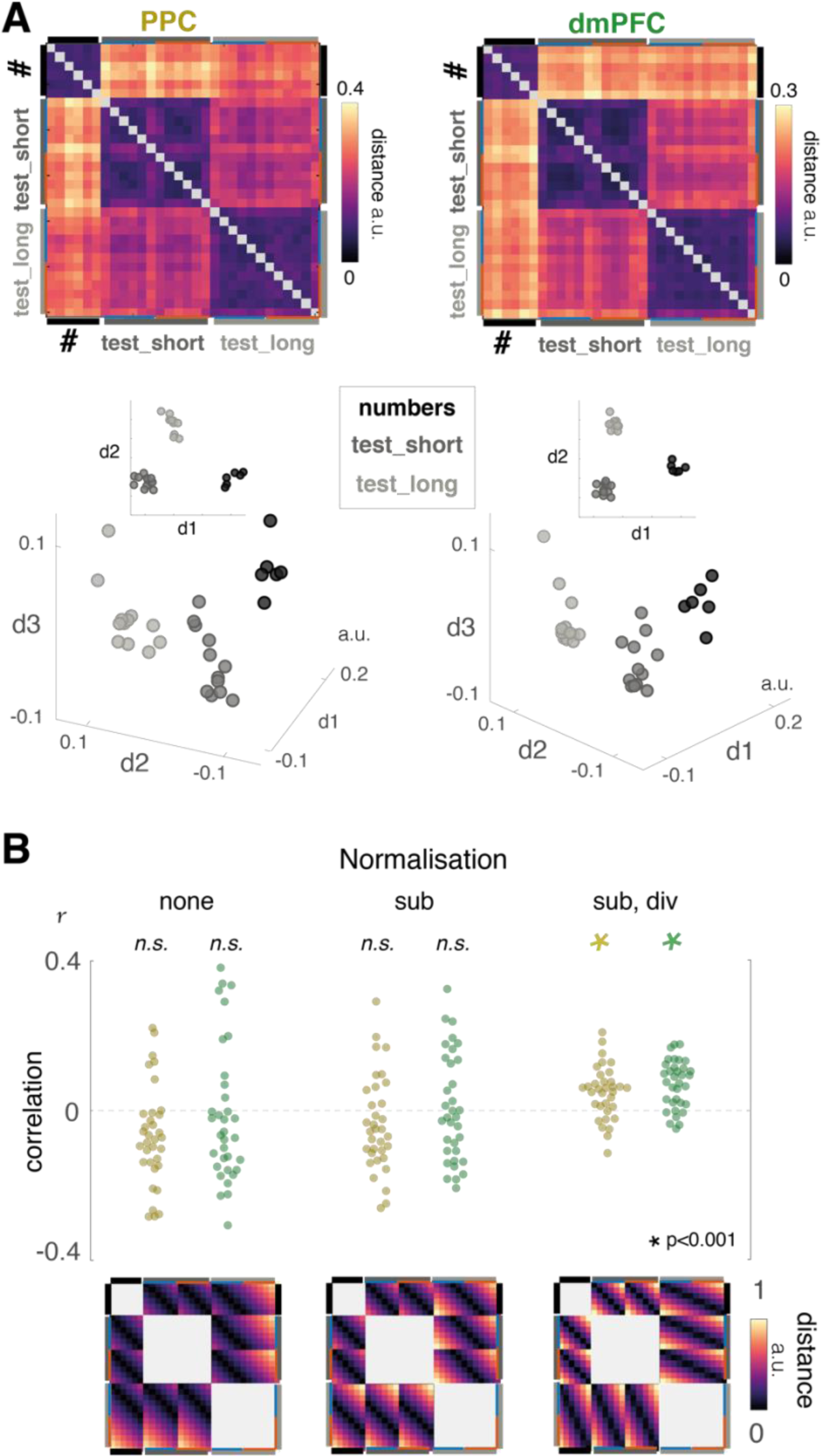
Cross task Normalization in Human fMRI. A. Cross-experiment Representational dissimilarity matrices (top) and multidimensional scaling plots (bottom) are plotted for the PPC (yellow) and dmPFC (green) ROIs reported in the main text. Each entry indicates the similarity of the neural patterns evoked by pairs of stimuli presented during the number localiser (black), test_short (dark grey) or test_long (light grey). We also overlay the context conditions (blue and red) in test_short and test_long for guidance. Note that as this analysis was intended to compare neural signals between the different experiment types, it used BOLD signals estimated using a GLM fit across all three experiments simultaneously. B. We constructed idealised distances matrices under different hypotheses of how this normalization could occur to quantify neural representations plotted in A. On the left, we devised a model of the ground-truth cross-experimental ordinal rankings (RDMnone). In the center, we considered a subtractive normalization scheme (RDMsub). Finally, on the right, we considered a subtractive and divisive normalization scheme (RDM_sub,div_). We considered only the distances between experiment types in this analysis, as indicated by the grey shaded regions of these idealized matrices. Pearson’s correlations between our observed and idealised matrices are plotted for PPC (yellow) and dmPFC (green), with stars indicating significance at p = 0.001.

**Supplementary Table 1:**
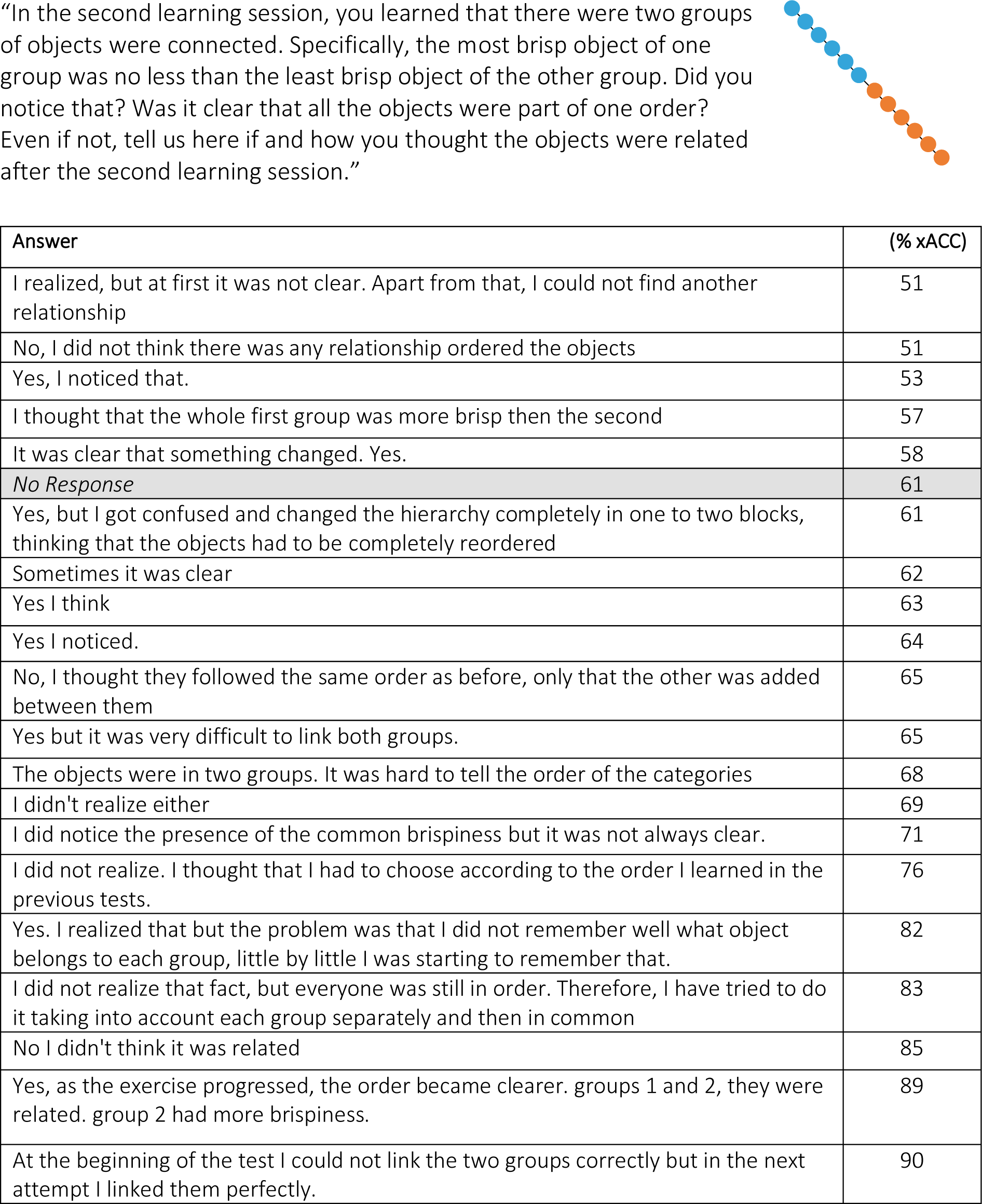
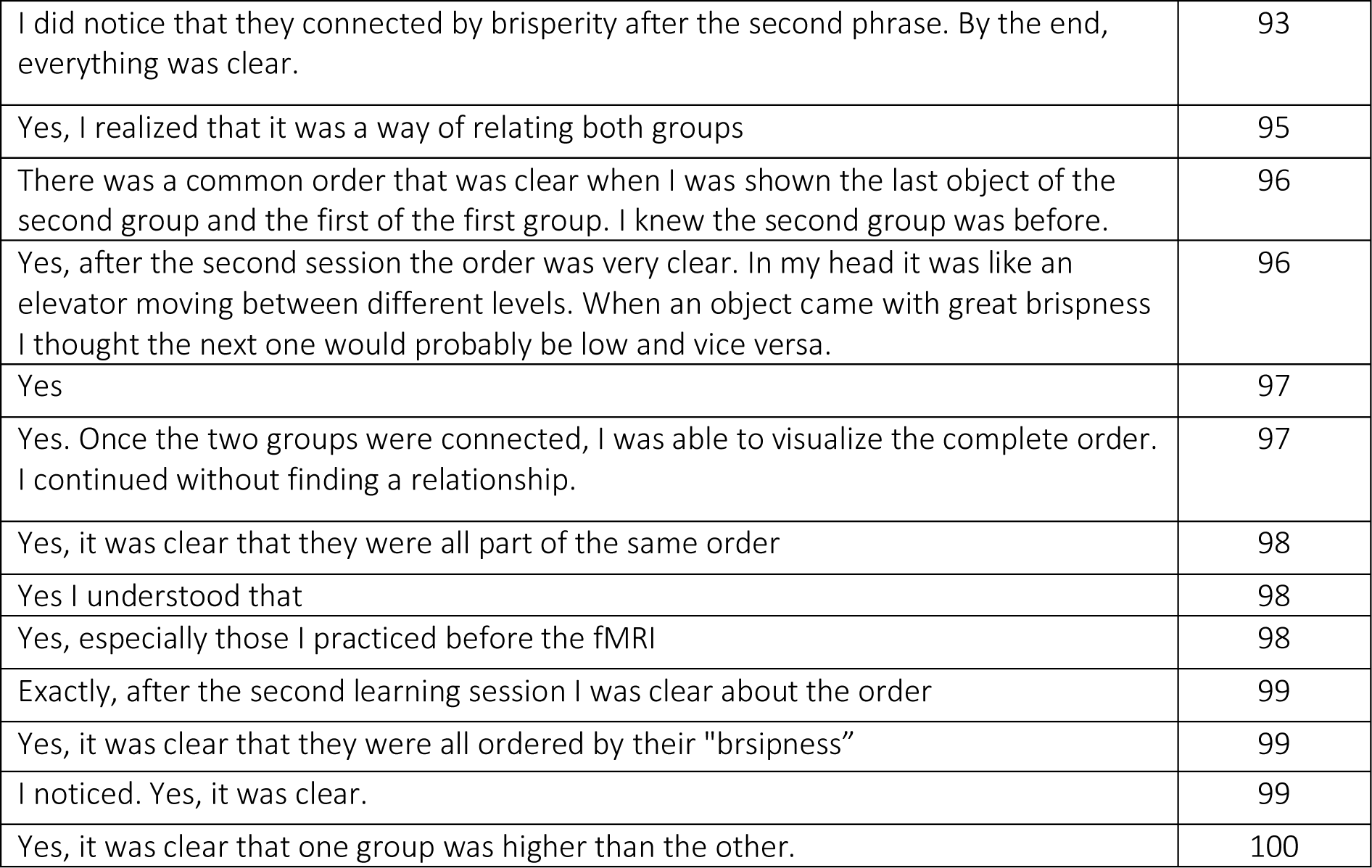
Participant responses to the debrief question below after completing the experiment. Answers were sorted by accuracy on comparisons between items initially in different sets (xAcc). Note that the debrief was voluntary, and one participant declined to answer.

| Answer | (% xACC) |
| --- | --- |
| I realized, but at first it was not clear. Apart from that, I could not find another relationship | 51 |
| No, I did not think there was any relationship ordered the objects | 51 |
| Yes, I noticed that. | 53 |
| I thought that the whole first group was more brisp then the second | 57 |
| It was clear that something changed. Yes. | 58 |
| *No Response* | 61 |
| Yes, but I got confused and changed the hierarchy completely in one to two blocks, thinking that the objects had to be completely reordered | 61 |
| Sometimes it was clear | 62 |
| Yes I think | 63 |
| Yes I noticed. | 64 |
| No, I thought they followed the same order as before, only that the other was added between them | 65 |
| Yes but it was very difficult to link both groups. | 65 |
| The objects were in two groups. It was hard to tell the order of the categories | 68 |
| I didn't realize either | 69 |
| I did notice the presence of the common brispiness but it was not always clear. | 71 |
| I did not realize. I thought that I had to choose according to the order I learned in the previous tests. | 76 |
| Yes. I realized that but the problem was that I did not remember well what object belongs to each group, little by little I was starting to remember that. | 82 |
| I did not realize that fact, but everyone was still in order. Therefore, I have tried to do it taking into account each group separately and then in common | 83 |
| No I didn't think it was related | 85 |
| Yes, as the exercise progressed, the order became clearer. groups 1 and 2, they were related. group 2 had more brispiness. | 89 |
| At the beginning of the test I could not link the two groups correctly but in the next attempt I linked them perfectly. | 90 |
| I did notice that they connected by brisperity after the second phrase. By the end, everything was clear. | 93 |
| Yes, I realized that it was a way of relating both groups | 95 |
| There was a common order that was clear when I was shown the last object of the second group and the first of the first group. I knew the second group was before. | 96 |
| Yes, after the second session the order was very clear. In my head it was like an elevator moving between different levels. When an object came with great brispness I thought the next one would probably be low and vice versa. | 96 |
| Yes | 97 |
| Yes. Once the two groups were connected, I was able to visualize the complete order. I continued without finding a relationship. | 97 |
| Yes, it was clear that they were all part of the same order | 98 |
| Yes I understood that | 98 |
| Yes, especially those I practiced before the fMRI | 98 |
| Exactly, after the second learning session I was clear about the order | 99 |
| Yes, it was clear that they were all ordered by their "brsipness" | 99 |
| I noticed. Yes, it was clear. | 99 |
| Yes, it was clear that one group was higher than the other. | 100 |

## References

1. Lake BM, Ullman TD, Tenenbaum JB, Gershman SJ. Building machines that learn and think like people. Behav Brain Sci. 2017;40: e253. doi:10.1017/S0140525X16001837

2. Morton NW, Preston AR. Concept formation as a computational cognitive process. Current Opinion in Behavioral Sciences. 2021;38: 83–89. doi:10.1016/j.cobeha.2020.12.005

3. Behrens TEJ, Muller TH, Whittington JCR, Mark S, Baram AB, Stachenfeld KL, et al. What Is a Cognitive Map? Organizing Knowledge for Flexible Behavior. Neuron. 2018;100: 490–509. doi:10.1016/j.neuron.2018.10.002

4. Lynn CW, Bassett DS. How humans learn and represent networks. Proc Natl Acad Sci USA. 2020;117: 29407–29415. doi:10.1073/pnas.1912328117

5. Tervo DGR, Tenenbaum JB, Gershman SJ. Toward the neural implementation of structure learning. Curr Opin Neurobiol. 2016;37: 99–105. doi:10.1016/j.conb.2016.01.014

6. Bellmund JLS, Gardenfors P, Moser EI, Doeller CF. Navigating cognition: Spatial codes for human thinking. Science. 2018;362. doi:10.1126/science.aat6766

7. Summerfield C, Luyckx F, Sheahan H. Structure learning and the posterior parietal cortex. Prog Neurobiol. 2019; 101717. doi:10.1016/j.pneurobio.2019.101717

8. Tolman EC. Cognitive maps in rats and men. Psychol Rev. 1948;55: 189–208.

9. Schapiro AC, Kustner LV, Turk-Browne NB. Shaping of object representations in the human medial temporal lobe based on temporal regularities. Curr Biol. 2012;22: 1622– 7. doi:10.1016/j.cub.2012.06.056

10. Schapiro AC, Rogers TT, Cordova NI, Turk-Browne NB, Botvinick MM. Neural representations of events arise from temporal community structure. Nat Neurosci. 2013;16: 486–92. doi:10.1038/nn.3331

11. Garvert MM, Dolan RJ, Behrens TE. A map of abstract relational knowledge in the human hippocampal-entorhinal cortex. Elife. 2017;6. doi:10.7554/eLife.17086

12. Zeithamova D, Preston AR. Temporal Proximity Promotes Integration of Overlapping Events. Journal of Cognitive Neuroscience. 2017;29: 1311–1323. doi:10.1162/jocn_a_01116

13. Whittington JCR, Muller TH, Mark S, Chen G, Barry C, Burgess N, et al. The Tolman-Eichenbaum Machine: Unifying Space and Relational Memory through Generalization in the Hippocampal Formation. Cell. 2020;183: 1249–1263.e23. doi:10.1016/j.cell.2020.10.024

14. Dordek Y, Soudry D, Meir R, Derdikman D. Extracting grid cell characteristics from place cell inputs using non-negative principal component analysis. eLife. 2016;5: e10094. doi:10.7554/eLife.10094

15. Klukas M, Lewis M, Fiete I. Efficient and flexible representation of higher-dimensional cognitive variables with grid cells. Bush D, editor. PLoS Comput Biol. 2020;16: e1007796. doi:10.1371/journal.pcbi.1007796

16. Saxe A, Nelli S, Summerfield C. If deep learning is the answer, what is the question? Nat Rev Neurosci. 2021;22: 55–67. doi:10.1038/s41583-020-00395-8

17. Lindsay G. Convolutional Neural Networks as a Model of the Visual System: Past, Present, and Future. J Cogn Neurosci. 2020; 1–15. doi:10.1162/jocn_a_01544

18. Barrett DGT, Hill F, Santoro A, Morcos AS, Lillicrap T. Measuring abstract reasoning in neural networks. arXiv:180704225 [cs, stat]. 2018 [cited 8 Oct 2020]. Available: http://arxiv.org/abs/1807.04225

19. Chang MB, Gupta A, Levine S, Griffiths TL. Automatically Composing Representation Transformations as a Means for Generalization. arXiv:180704640 [cs, stat]. 2019 [cited 5 May 2021]. Available: http://arxiv.org/abs/1807.04640

20. Lake BM, Salakhutdinov R, Tenenbaum JB. Human-level concept learning through probabilistic program induction. Science. 2015;350: 1332–8. doi:10.1126/science.aab3050

21. Higgins I, Sonnerat N, Matthey L, Pal A, Burgess CP, Bosnjak M, et al. SCAN: Learning Hierarchical Compositional Visual Concepts. arXiv:170703389. 2017.

22. Morton NW, Sherrill KR, Preston AR. Memory integration constructs maps of space, time, and concepts. Current Opinion in Behavioral Sciences. 2017;17: 161–168. doi:10.1016/j.cobeha.2017.08.007

23. Bernardi S, Benna MK, Rigotti M, Munuera J, Fusi S, Salzman CD. The Geometry of Abstraction in the Hippocampus and Prefrontal Cortex. Cell. 2020; S0092867420312289. doi:10.1016/j.cell.2020.09.031

24. Sheahan H, Luyckx F, Nelli S, Teupe C, Summerfield C. Neural state space alignment for magnitude generalization in humans and recurrent networks. Neuron. 2021;109: 1214–1226.e8. doi:10.1016/j.neuron.2021.02.004

25. Luyckx F, Nili H, Spitzer B, Summerfield C. Neural structure mapping in human probabilistic reward learning. Elife. 2019;8. doi:10.7554/eLife.42816

26. Woocher FD, Glass AL, Holyoak KJ. Positional discriminability in linear orderings. Memory & Cognition. 1978;6: 165–173. doi:10.3758/BF03197442

27. Treichler FR, Van Tilburg D. Concurrent conditional discrimination tests of transitive inference by macaque monkeys: List linking. Journal of Experimental Psychology: Animal Behavior Processes. 1996;22: 105–117. doi:10.1037/0097-7403.22.1.105

28. Horst JS, Hout MC. The Novel Object and Unusual Name (NOUN) Database: A collection of novel images for use in experimental research. Behav Res. 2016;48: 1393–1409. doi:10.3758/s13428-015-0647-3

29. D’Amato MR, Colombo M. The symbolic distance effect in monkeys (Cebus apella). Animal Learning & Behavior. 1990;18: 133–140. doi:10.3758/BF03205250

30. Kumaran D, McClelland JL. Generalization through the recurrent interaction of episodic memories: a model of the hippocampal system. Psychol Rev. 2012;119: 573–616. doi:10.1037/a0028681

31. Chen S, Swartz KB, Terrace HS. Knowledge of the Ordinal Position of List Items in Rhesus Monkeys. Psychol Sci. 1997;8: 80–86. doi:10.1111/j.1467-9280.1997.tb00687.x

32. Flesch T, Juechems K, Dumbalska T, Saxe A, Summerfield C. Rich and lazy learning of task representations in brains and neural networks. Neuroscience; 2021 Apr. doi:10.1101/2021.04.23.441128

33. Okazawa G, Hatch CE, Mancoo A, Machens CK, Kiani R. Representational geometry of perceptual decisions in the monkey parietal cortex. Cell. 2021; S0092867421006528. doi:10.1016/j.cell.2021.05.022

34. McClelland JL, McNaughton BL, O’Reilly RC. Why there are complementary learning systems in the hippocampus and neocortex: insights from the successes and failures of connectionist models of learning and memory. Psychol Rev. 1995;102: 419–57.

35. Kumaran D, Hassabis D, McClelland JL. What Learning Systems do Intelligent Agents Need? Complementary Learning Systems Theory Updated. Trends Cogn Sci. 2016;20: 512–534. doi:10.1016/j.tics.2016.05.004

36. Hunt LT, Daw ND, Kaanders P, MacIver MA, Mugan U, Procyk E, et al. Formalizing planning and information search in naturalistic decision-making. Nat Neurosci. 2021 [cited 2 Jul 2021]. doi:10.1038/s41593-021-00866-w

37. Liu Y, Dolan RJ, Kurth-Nelson Z, Behrens TEJ. Human Replay Spontaneously Reorganizes Experience. Cell. 2019;178: 640–652 e14. doi:10.1016/j.cell.2019.06.012

38. Zenke F, Poole B, Ganguli S. Continual Learning Through Synaptic Intelligence. arXiv:170304200. 2017.

39. Kirkpatrick J, Pascanu R, Rabinowitz N, Veness J, Desjardins G, Rusu AA, et al. Overcoming catastrophic forgetting in neural networks. Proc Natl Acad Sci U S A. 2017;114: 3521–3526. doi:10.1073/pnas.1611835114

40. Kurth-Nelson Z, Economides M, Dolan RJ, Dayan P. Fast Sequences of Non-spatial State Representations in Humans. Neuron. 2016;91: 194–204. doi:10.1016/j.neuron.2016.05.028

41. Wimmer GE, Liu Y, Vehar N, Behrens TEJ, Dolan RJ. Episodic memory retrieval success is associated with rapid replay of episode content. Nat Neurosci. 2020;23: 1025–1033. doi:10.1038/s41593-020-0649-z

42. Nour MM, Liu Y, Arumuham A, Kurth-Nelson Z, Dolan RJ. Impaired neural replay of inferred relationships in schizophrenia. Cell. 2021; S0092867421007479. doi:10.1016/j.cell.2021.06.012

43. Wimmer GE, Daw ND, Shohamy D. Generalization of value in reinforcement learning by humans. The European journal of neuroscience. 2012;35: 1092–104. doi:10.1111/j.1460-9568.2012.08017.x

44. Botvinick MM, Braver TS, Barch DM, Carter CS, Cohen JD. Conflict monitoring and cognitive control. Psychol Rev. 2001/08/08 ed. 2001;108: 624–52.

45. Hubbard EM, Piazza M, Pinel P, Dehaene S. Interactions between number and space in parietal cortex. Nat Rev Neurosci. 2005;6: 435–48. doi:10.1038/nrn1684

46. Walsh V. A theory of magnitude: common cortical metrics of time, space and quantity. Trends Cogn Sci. 2003;7: 483–8. doi:10.1016/j.tics.2003.09.002

47. Yu LQ, Park SA, Sweigart SC, Boorman ED, Nassar MR. Do grid codes afford generalization and flexible decision-making? arXiv:210616219 [q-bio]. 2021 [cited 16 Sep 2021]. Available: http://arxiv.org/abs/2106.16219

48. Niv Y, Daniel R, Geana A, Gershman SJ, Leong YC, Radulescu A, et al. Reinforcement Learning in Multidimensional Environments Relies on Attention Mechanisms. Journal of Neuroscience. 2015;35: 8145–8157. doi:10.1523/JNEUROSCI.2978-14.2015

